# The regulatory architecture of gene expression variation in *C. elegans* revealed by multi-strain allele-specific analysis

**DOI:** 10.1101/2024.10.15.618466

**Authors:** Avery Davis Bell, Francisco Valencia, Annalise B. Paaby

## Abstract

An outstanding question in the evolution of gene expression is the composition of the underlying regulatory architecture and the processes that shape it. Mutations affecting a gene’s expression may reside locally in *cis* or distally in *trans*; the accumulation of these changes, their interactions, and their modes of inheritance influence how traits are expressed and how they evolve. Here, we interrogated gene expression variation in *C. elegans*, including the first allele-specific expression analysis in this system, capturing effects in *cis* and in *trans* that govern gene expression differences between the reference strain N2 and seven wild strains. We observed extensive compensatory regulation, in which opposite effects in *cis* and *trans* at individual genes mitigate expression differences among strains, and that genes with expression differences exhibit strain specificity. As the genomic distance increased between N2 and each wild strain, the number of genes with expression differences also increased. We also report for the first time that expression-variable genes are lower expressed on average than genes without expression differences, a trend that may extend to humans and *Drosophila melanogaster* and may reflect the selection constraints that govern the universal anticorrelation between gene expression and rate of protein evolution. Together, these and other observed trends support the conclusion that many *C. elegans* genes are under stabilizing selection for expression level, but we also highlight outliers that may be biologically significant. To provide community access to our data, we introduce an easily accessible, interactive web application for gene-based queries: https://wildworm.biosci.gatech.edu/ase/.

**Author Summary:** Genes are first expressed as RNAs, which act as templates in the synthesis of proteins, the workhorse molecules of the cell. Naturally-occurring mutations can influence how RNA expression occurs; to evaluate how RNA expression varies among individuals and how such patterns evolve, the authors measured expression in wild strains of the nematode *C. elegans* and in offspring derived from crosses between them. This study describes the architecture of RNA expression variation in this essential model organism and reports novel patterns that may extend to insects and humans.

## Introduction

Gene expression is an essential step in the translation of genotype to phenotype. Thus, to understand the evolution of traits and genomes, we need to elucidate the regulatory mechanisms, modes of inheritance, and biological processes shaping gene expression variation within a system. Mutational variants that affect gene expression may act in *cis,* locally within the focal gene haplotype, such as promoter variants; or in *trans*, on a separate molecule and potentially affecting all targets, such as mutations in transcription factors (Signor and Nuzhdin 2018). Regulatory variants that mediate gene expression may represent adaptive change, neutral differences, or relaxed selection (reviewed in, *e.g.,* Landry *et al*. 2007b; Fay and Wittkopp 2008; Romero *et al*. 2012; Signor and Nuzhdin 2018, 2019; Price *et al*. 2022a; Hill *et al*. 2020). They may also act to stabilize expression by dampening changes to expression induced by other variants.

An incisive way to study gene expression regulation is to examine variation by simultaneously capturing expression among wild strains and their F1 hybrid offspring (Wittkopp *et al*. 2004; Landry *et al*. 2007a). Within the F1, expression differences observed between the parental alleles (allele-specific expression, ASE) may be assigned to mutations in *cis*, on the same molecule, because the diffusible *trans* environment is shared within cells (Yan *et al*. 2002; Cowles *et al*. 2002). Thus, comparisons of expression between alleles, between parents, and between F1s and parents enable inference of the regulatory architecture and inheritance mode of gene expression (Wittkopp *et al*. 2004; McManus *et al*. 2010). This approach has been employed in a number of systems to interrogate various phenomena, including domestication, adaptation, and speciation in wild and crop plants (Bao *et al*. 2019; He *et al*. 2016; He *et al*. 2012; Lemmon *et al*. 2014; Rhone *et al*. 2017; Steige *et al*. 2017; Steige *et al*. 2015; Verta *et al*. 2016; Zhang and Borevitz 2009); adaptation and the evolution of embryogenesis in *Drosophila* (Cartwright and Lott 2020; Juneja *et al*. 2016; Coolon *et al*. 2014; McManus *et al*. 2010); speciation and *cis* regulatory variation in mice (Crowley *et al*. 2015; Mack *et al*. 2016); human-specific regulatory evolution in chimpanzee-human hybrid cell lines (Gokhman *et al*. 2021; Starr *et al*. 2023; Wang *et al*. 2024); RNA and protein regulation in yeast (Artieri and Fraser 2014; Muzzey *et al*. 2014; Wang *et al*. 2015); and speciation and evolution of reproductive mode in nematodes (Sanchez-Ramirez *et al*. 2021; Xie *et al*. 2022; Viswanath and Cutter 2023).

*C. elegans* has long been a leading developmental and genetic model organism (Sternberg *et al*. 2024), and the recent establishment of a global collection of wild strains has pushed *C. elegans* to the forefront of quantitative genetics and evolutionary genomics research (Frézal and Félix 2015; Andersen and Rockman 2022; Crombie *et al*. 2024; Crombie *et al*. 2019; Cook *et al*. 2017). Yet, while the genetic basis of expression variation has been interrogated via well-powered eQTL studies (Rockman *et al*. 2010; Vinuela *et al*. 2010; Francesconi and Lehner 2014; Kamkina *et al*. 2016; Evans and Andersen 2020; Zhang *et al*. 2022), the regulatory architecture and inheritance mode of gene expression variation in *C. elegans* has not been assessed by allele-specific analyses. However, the biology of *C. elegans* offers rich opportunity for investigating gene expression variation and its evolution, beyond its well-established resources. *C. elegans* strains persist as predominantly selfing lineages in diverse ecological habitats across the globe, with the population structure exhibiting a spectrum of genetic divergence between wild strains, from closely related to highly diverged (Barriere and Felix 2005b; Barriere and Felix 2005a; Crombie *et al*. 2024; Crombie *et al*. 2019; Lee *et al*. 2021). The genomes harbor extensive linkage disequilibrium, including long haplotypes arising from historical adaptive sweeps, and inter-strain crosses often exhibit fitness deficits, suggesting disruption of the selfed, co-adapted genotype combinations (Barriere and Felix 2005a; Dolgin *et al*. 2007; Rockman and Kruglyak 2009; Andersen *et al*. 2012). Thus, *C. elegans* is optimally suited to facilitate investigations into whether and how genetic divergence translates to differences in expression, into the scope and correlates of compensatory interactions in the evolution of gene expression regulation, and into the broader evolutionary pressures shaping these trends.

The role of compensatory interactions in the evolution of gene expression is incompletely understood, but a growing body of literature suggests that such dynamics are influential and pervasive. Gene expression changes often fail to result in protein-level changes (Schrimpf *et al*. 2009; Khan *et al*. 2013; Brion *et al*. 2020; Buccitelli and Selbach 2020) and regulatory changes to expression arising in *cis* often fail to produce overall differences in gene expression, implying that they are compensated by regulation in *trans* (Landry *et al*. 2005; Signor and Nuzhdin 2018, 2019). Studies have reported such compensation of *cis-*regulated differences in hybrids of different species, subspecies, and occasionally strains of fruit flies, sticklebacks, cotton, mice, yeast, spruce, and more (Landry *et al*. 2005; Goncalves *et al*. 2012; Bao *et al*. 2019; Coolon *et al*. 2014; McManus *et al*. 2010; Metzger *et al*. 2017; Verta and Jones 2019; Verta *et al*. 2016; Signor and Nuzhdin 2018, 2019). In yeast, dissection of the *trans* regulatory architecture of a single gene estimated hundreds of variants mediating its expression, with extensive compensatory interactions and evidence of stabilizing selection on overall expression level (Metzger and Wittkopp 2019). However, methodological constraints and analytical artifacts can limit confidence at both protein and RNA levels, hindering widespread inferences (Price *et al*. 2022a; Fraser 2022; Price *et al*. 2022b; Fraser 2019; Buccitelli and Selbach 2020). In *C. elegans*, fitness-related traits exhibit compensatory-like architecture, with epistasis and tightly-linked opposite-direction effects shaping fertility and fecundity (Noble *et al*. 2017; Bernstein *et al*. 2019); these traits may be governed by similar dynamics at the level of gene expression. Overall, the extent to which *C. elegans* gene expression has evolved compensatory interactions remains an open question.

Here, to elucidate the regulatory architecture and evolutionary dynamics shaping gene expression in *C. elegans*, we examine expression variation in seven intraspecific crosses. In each, the reference strain N2 was crossed to one of seven wild strains representing a spectrum of genomic differentiation from the reference. We define the regulatory patterns and inheritance modes of expression variation in this system, then assess how regulatory effects are influenced by factors such as nucleotide diversity, genome evolutionary history, gene essentiality and biological role, and expression level.

## Materials and methods

### Experimental methods

A detailed protocol describing the experimental methods is available at protocols.io (dx.doi.org/10.17504/protocols.io.5jyl8p15rg2w/v1, Bell *et al*. 2024).

#### Worm strains

**Table S1** provides the complete list of strains used in this study. In selecting parental strains to cross with the N2 laboratory reference strain to generate F1s in which to investigate allele-specific expression (ASE), we aimed to represent the range of nucleotide diversity present in the species as well as capture outlier strains. All chosen strains differed at more than 127,000 nucleotides from N2 (>1.27 variants per kilobase average) (per CaeNDR, Crombie *et al*. 2024) to ensure that the F1s harbored many genes with differences from the reference in coding regions. To ensure that we generated F1s with one copy of the genome from each parent, rather than N2 self-progeny, we used the N2 strain feminized via a deletion of *fog-2* as the N2 ‘female’ parent (referred to in the text as N2*^fog-2^*, strain CB4108): *fog-2* deficient hermaphrodites are incapable of producing sperm and therefore function as females (Hodgkin 2002; PedersenSchedl and Kimble 1988).

#### Worm husbandry

We thawed fresh aliquots of each wild strain and grew them without starving for at least three generations, but for no more than one month, prior to starting the experiment. We followed standard protocol (Stiernagle 2006) for worm culture, using 1.25% agarose plates to prevent wild strains’ burrowing. Prior to the start of the experiment, all strains were maintained at 18°C to allow slower growth of large quantities of worms and to avoid inducing QX1211’s mortal germline phenotype, which is more penetrant at higher temperatures (Frezal *et al*. 2018).

#### Generating parallel F1 crosses and self-progeny

As detailed in our protocol (Bell *et al*. 2024), we first bleach synchronized all parental strains to ensure that the parents that would be mated were of similar developmental stage, as parental age can impact offspring development and transcriptional program (Perez *et al*. 2017; Webster *et al*. 2023). To ensure that we would have many L4 parent worms to move to mating plates, we grew several plates of all bleached strains at 18°C, 19°C, and 20°C, and additionally grew the N2*^fog-2^* parent (from whom we needed the highest number of worms) at room temperature.

After allowing these worms to grow for two days, we generated mating plates by placing 60-80 N2*^fog-2^* L4 pseudo-hermaphrodites onto each of five 6cm plates with small bacteria spots and added 40 L4 males of the appropriate strain to each plate. We concurrently moved 80 individual L4 hermaphrodites to each of three 6cm plates for each parental strain (N2 and seven wild strains) to simultaneously generate the parental strains used for sequencing from self-matings while the F1 crosses were generated from cross-matings.

After allowing mating for 48 hours, we collected and synchronized the offspring for the crosses and self-matings by collecting all parental worms and embryos from the bacterial lawn, treating with bleach, and allowing embryos to develop into L1 larvae and arrest over 30 hours in liquid buffer. After 30 hours, L1s were transferred directly to the bacterial lawn of 6cm plates at a density of ∼400 L1s per plate.

After allowing the worms to develop for ∼36 hours, we removed males from the F1 plates as soon as they were detectable and screened the parental plates for any spontaneously generated males, which were also removed. Plates used for RNA sequencing (at least 3 per strain) had all males removed as L4s or young adults.

#### Worm harvesting

Worms were harvested as day 1 reproductively mature young adults: when most worms were gravid with embryos and laid embryos were visible on the plates. Because developmental timing differs across wild strains (Gems and Riddle 2000; Stastna *et al*. 2015; Zhang *et al*. 2021; Hodgkin and Doniach 1997; Poullet *et al*. 2015; Harvey and Viney 2007), we chose to match developmental stage rather than hours of development; even so, all worms reached reproductive maturity and were harvested within three hours of each other. Worms were rinsed off plates, washed with M9 buffer, and resuspended in TRIzol (Invitrogen #15596026) in 3 tubes (replicates) per strain before immediate flash freezing in liquid nitrogen and storage at −80°C until RNA extraction.

#### RNA library preparation and sequencing

RNA was extracted from worms stored in TRIzol (Invitrogen #15596026) following standard procedure (following He 2011, see Bell *et al*. 2024) using a TRIzol (Invitrogen #15596026) chloroform (Fisher #C298-500) extraction and RNeasy columns (Qiagen #74104). This extraction was performed in 3 batches of 15 over two consecutive days, with one replicate from each strain included in each batch. RNA was stored at −80°C for ∼1 week prior to library generation. Library preparation and sequencing for all samples was performed by the Molecular Evolution Core Laboratory at the Georgia Institute of Technology. Following RNA quality checks (all RINs 9.8 or greater), mRNA was enriched from 1μg RNA with the NEBNext Poly(A) mRNA magnetic isolation module (NEB #E7490) and sequencing libraries generated using the NEBNext Ultra II directional RNA library preparation kit (NEB #E7760) with 8 cycles of PCR. Libraries were quality checked and fluorometrically quantified prior to pooling and sequencing. Libraries were sequenced on an Illumina NovaSeq X using a 300 cycle 10B flowcell. A median of 65 million 150x150bp sequencing read pairs were generated per library (range 25-93 million, **Table S1**).

### Analytical methods

The code written for this study is available at https://github.com/paabylab/wormase.

#### Expression quantification

Before expression quantification, we generated strain-specific transcriptomes as described previously (Bell *et al*. 2023) by inserting known SNV and INDEL polymorphisms (from the CeNDR (Cook *et al*. 2017; Crombie *et al*. 2024) 2021021 release hard-filter VCF) into the *C. elegans* reference genome (ws276 from WormBase, Sternberg *et al*. 2024) and extracting transcripts. We created pseudo-diploid strain transcriptomes by combining these strain-specific transcriptomes for the two parent strains. Tools used in generating these transcriptomes included g2gtools (v0.1.31) (https://github.com/churchill-lab/g2gtools), gffread (v0.12.7) (Pertea and Pertea 2020), seqkit (v0.16.1) (Shen *et al*. 2016), and bioawk (v1.0) (https://github.com/lh3/bioawk). For comparison purposes, we also created pseudo-diploid and strain-specific transcriptomes using script *create_personalized_transcriptome.py* from the Ornaments code suite (initial version) (Adduri and Kim 2024) tool, with the ws286 genome build and 20220216 CeNDR VCF.

For quantification used in allele-specific expression and differential expression analyses, we estimated allele-specific and total RNA counts using EMASE (emase-zero v0.3.1) (Raghupathy *et al*. 2018) with input quantifications generated by running Salmon (v1.4) (Patro *et al*. 2017) against the pseudo-diploid transcriptomes. Specifically, we generated a salmon index for the diploid transcriptome using *salmon index* with options *-k 31 --keepDuplicates* (no decoy, all other parameters default). To prepare RNA-seq data for quantification, we trimmed Illumina adapters using trimmomatic (v0.39) (Bolger *et al*. 2014) with parameters *ILLUMINACLIP: TruSeq3-PE-2.fa:1:30:12:2:True*. Salmon quantification with equivalence class outputs saved was performed against the pseudo-diploid transcript’s index with *salmon quant -l ISR --dumpeq --fldMean <*sample-specific mean> *--fldSD* <sample-specific SD> *--rangeFactorizationBins 4 -- seqBias --gcBias*. Salmon outputs were converted to .bin inputs for emase-zero using *alntools salmon2ec* (v0.1.1) (https://churchill-lab.github.io/alntools/). Finally, emase-zero was run on this input using parameters *--model 4 -t 0.0001 -i 999.* For comparison, we separately generated quantification estimates using kallisto (v0.50.1) (Bray *et al*. 2016) against strain-specific transcriptomes generated by Ornaments, and estimated allele-specific RNA counts using *ornaments quant* (initial version), which implements WASP (van de Geijn *et al*. 2015)-style allele-specific quantification on top of kallisto quantification and includes INDELs in its analysis. Workflows to perform these steps are in our code repository in internal directories: *data_generation_scripts/getdiploidtranscriptomes; data_generation_scripts/emase; data_generation_scripts/ornaments*

We used DESeq2 (v1.42.0) (Love *et al*. 2014) to obtain final RNA quantifications for downstream modeling. For differential expression analyses, we used the “total” column of the “gene.counts” output from emase-zero. For allele-specific analyses, we used the allelic counts columns of the “gene.counts” output from emase-zero. For kallisto quantifications, transcript TPMs were combined to gene-level, normalized quantifications for DESeq2 using tximport (v1.30.0) (Soneson *et al*. 2015). In all cases, genes with at least 10 total reads summed across samples were retained for downstream analysis. For obtaining general best expression quantification estimates (rather than for differential expression modeling), we used DESeq2’s variance stabilizing transformation (*vst* function) to get log-scale, variance normalized, length and library size normalized gene expression estimates.

#### Age estimation

We estimated each sample’s age in hours against a developmental timing ‘ruler’ from the N2 strain via RAPToR (v1.2.0) (Bulteau and Francesconi 2022) using DESeq2’s *vst* corrected gene counts from total emase-zero outputs. The age reference used (provided with RAPToR) was *Cel_YA_2*. The script used to perform this analysis is in our code repository: *data_classification_scripts/RAPToR.R*

#### Differential expression and allele-specific expression calling

Each sample was assigned to its generation-strain group (*e.g.,* CB4856 F1). Total gene counts from emase-zero “total” gene.counts output were binomially negatively modeled by DESeq2 as

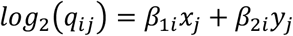

Where, for gene *i*, sample *j*, *q* is proportional to RNA concentration/counts (Love *et al*. 2014), βs give the effects for gene *i* for RNA extraction replicate (*x*) and each generation-strain pair (*y*). The Wald test was used for significance testing. Results were pulled out for each pairwise comparison of interest using DESeq2’s contrasts. All log_2_ fold changes were adjusted using *ashr* (v2.2-63) (Stephens 2016). For differential expression to be called, both a fold change of greater than 1.5 after *ashr* adjustment (for significance testing and calling) and a genome-wide adjusted *p* value less than 0.05 were required.

For genes to be included in allele-specific expression analyses, we required them to have 5 gene and allele-specific alignments. The total counts of alignments per gene and those that were gene and allele-specific were derived by analyzing of salmon’s equivalence class output file, which assigns equivalence classes of kmers to transcripts from which they derive and gives the counts of reads aligning to each equivalence class. We investigated several thresholds of gene- and allele-specific alignments for considering a gene ASE-informative; we found that our RNA sequencing was deep enough that once genes in each F1 genotype had more than three allele- and gene-specific alignments in each sample from that genotype, they usually had many allele- and gene-specific alignments. Therefore, we required genes to have a slightly conservative five allele- and gene-specific alignments to be considered ASE-informative.

To model allele-specific expression in the F1s, each allele’s count was represented in its own column in the model matrix. Within each strain, each sample was assigned its sample blocking factor, controlling for sample during modeling. We used DESeq2’s negative binomial modeling to model allele counts:

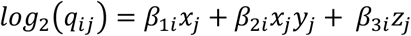

Where, for gene *i*, allele (rather than sample) *j*, *q* is proportional to allelic RNA concentration/counts (Love *et al*. 2014), β*_1_* gives the effect of RNA extraction replicate (*x*), β*_2_* gives the effect of the interaction between RNA extraction replicate and specific sample (*xy*), and β*_3_* gives the effect of the allele/genotype (*z*). Library size correction was not used in this modeling because all comparisons were being done within-sample, where library size was identical, and counts were of alleles rather than total. Library size correction was excluded by setting all DESeq2 size factors to 1 prior to differential expression testing. Results were extracted for each allelic pairwise comparison of interest and were used in downstream analysis for ASE-informative genes. ASE-informative genes were considered to have ASE if their *ashr*- adjusted fold change was greater in magnitude than 1.5 (equivalent to having 60% of alleles come from one haplotype) and their genome-wide-adjusted *p* value was less than 0.05 (the same thresholds required for DE calls; fold change threshold used in both significance testing and calling). Both log_2_ fold changes and the proportion of alleles deriving from the reference and alternate genomes were used for downstream analytical interpretation; alternate allele proportion was calculated from the *ashr*-adjusted log_2_ fold change (*LFC)* as

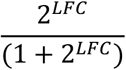

The scripts used for these analyses are in our code repository: equivalence class processing for ASE-informative decisions in *data_generation_scripts/ salmonalleleeqclasses.py*; ASE and DE modeling in *data_classification_scripts/ ase_de_annotategenes_deseq2_fromemaseout.R*

#### Inheritance mode classifications

Inheritance mode categories were called from differential expression testing results (from global RNA counts); categories and definitions followed McManus *et al*. (2010) and others, with thresholds tuned for our specific statistical testing framework. All *p* values were genome-wide adjusted and FCs/LFCs (fold changes/log_2_ fold changes) were *ashr* adjusted. Genes were called ‘no_change’ if there was no DE between the parents, between the F1 and the N2 parent, or between the F1 and the other parent (all *p* > 0.05 or |FC| < 1.5). Genes were called ‘overdominant’ if the F1 had higher expression than both parents (FC > 1.5 and *p* < 0.05). Genes were called ‘underdominant’ if the F1 had lower expression than both parents (FC < −1.5 and *p* < 0.05). Genes were called ‘N2_dominant’ if the parents were differentially expressed and the F1 was potentially differentially expressed from the wild parent in the same direction as N2 was (N2 *vs* wild strain |FC| > 1.5 and *p* < 0.05, F1 *vs* wild strain *p* < 0.05 and FC in the same direction as N2’s), or if the parents were potentially differentially expressed and the F1 was differentially expressed in the same direction from the wild parent as N2 was (N2 *vs* wild strain *p* < 0.05 and FC in the same direction as F1’s; F1 *vs* wild strain |FC| > 1.5 and *p* < 0.05). Genes were called ‘alt_dominant’ the same way as *N2_dominant* but requiring the F1 to be differentially expressed from the N2 parent in the same way as its wild parent. Genes were called ‘additive’ if the parent strains were differentially expressed (*p* < 0.05 and |FC| > 1.5) and the F1 had nominally called differential expression with expression amount falling between the two parents (*p* < 0.05, FC > 0 if parental FC > 0 and FC < 0 if parental FC < 0). Therefore, genes with midparental gene expression closer to the expression level of one parent or another, which might more formally be termed ‘nearly additive,’ are included in the ‘additive’ category. Genes whose DE results did not meet any of the above requirements were called ‘ambiguous’, for example when parental DE was not called but the F1 had DE called from one parent (these genes might be either additively inherited or dominantly inherited, but the statistical evidence was not strong enough to discern which). The inheritance mode classification script is available in our code repository: *data_classification_scripts/ f1_parental_inhmode_withinstrain.R*

#### Regulatory pattern and related classifications

Regulatory pattern categories were called from comparisons of allele-specific expression (N2 vs. wild strain allele) calls and differential expression (N2 vs. wild strain total RNA counts) calls; categories and definitions followed McManus *et al*. (2010) and others, with the specific thresholds tuned for our specific statistical testing framework. All *p* values were genome-wide adjusted and FCs/LFCs (fold change/log_2_ fold changes) were *ashr* adjusted and categorizations were only considered if genes were ASE-informative. Genes were called ‘conserved’ if they had neither ASE nor DE (both allelic and strain-wise *p* > 0.05 and |FC| < 1.5). Genes were called ‘cis’ *(i.e., cis*-only or *cis-*dominant regulatory divergence) if ASE and DE were both present and in the same direction and if their 99.9% confidence intervals on effect size overlapped (allelic *p* < 0.05 and |FC| > 1.5, strain-wise *p* < 0.05 without FC threshold, log2FC(DE) / log2FC(ASE) > 0). Genes were called ‘trans’ (*i.e., trans*-only or *trans-*dominant regulatory divergence) if they did not have ASE but did have DE (allelic *p* > 0.05, strain-wise *p* < 0.05 and |FC| > 1.5). Genes were called ‘enhancing’ (i.e. *cis-trans* enhancing or *cis+trans*) if they had both ASE and DE in the same direction and DE was of greater magnitude than ASE with non-overlapping 99.9% confidence intervals of the ASE and DE estimates (ASE *p* < 0.05 and |FC| > 1.5 and DE *p* < 1, or ASE *p* < 0.05 and DE *p* < 0.05 and |FC| > 1.5; and log2FC(DE) / log2FC(ASE) > 1). Genes were called ‘compensating’ (*i.e. cis* and *trans* regulatory changes in opposite directions, with the *cis* effect larger than the *trans* effect) if they had ASE and DE in the same direction with larger ASE than DE and non-overlapping 99.9% confidence intervals on the ASE and DE estimates (0 > log2FC(DE)/log2FC(ASE) > 1, allelic *p* < 0.05 and |FC| > 1.5 and strain-wise *p* < 0.05 *or* allelic *p* < 0.05 and strain-wise *p* < 0.05 and |FC| > 1.5). Genes were called ‘compensatory’ (*i.e.*, *cis* and *trans* regulatory changes in opposite directions, with *trans* changes fully offsetting the *cis* changes) if there was ASE but not DE (allelic *p* < 0.05 and |FC| > 1.5, strain-wise *p* > 0.05). Genes were called ‘overcompensating’ (*i.e., cis* and *trans* regulatory changes in opposite directions, with the *trans* change more than offsetting the *cis* effect) if they had ASE and DE in different directions with non-overlapping 99.9% confidence intervals on the ASE and DE estimates (log2FC(DE)/log2FC(ASE) < 0; allelic *p* < 0.05 and |FC| > 1.5 and strain-wise *p* < 0.05 *or* allelic *p* < 0.05 and strain-wise *p* < 0.05 and |FC| > 1.5). Genes were called ‘ambiguous’ if they did not meet the above criteria, *e.g.,* when ASE and DE were called with overlapping widened confidence intervals on opposite-direction effects. The regulatory pattern classification script is in our code repository: *data_analysis_scripts/ase_de_cistransclassifications.R*

We simplified these regulatory patterns for ease of understanding and visualization in a couple of ways. First, genes were classified as ‘cis-trans opposing’ anytime they had opposite direction *cis* and *trans* effects, *i.e.,* when their regulatory pattern was ‘compensating’*, ‘*compensatory’, or ‘over-compensating’. Second, we used the regulatory patterns to investigate compensation in a more targeted way, classifying genes as compensated if their simplified regulatory pattern was ‘cis-trans opposing’ and as not compensated if their regulatory pattern was ‘cis’ or ‘enhancing’. Genes without *cis* regulatory changes therefore are neither compensated or not compensated and were not included in compensation-specific analyses.

To evaluate the extent of compensatory regulatory effects, *i.e.*, to assess whether individual genes showed an enrichment for effects in both *cis* and *trans*, and specifically for opposing effects in *cis* versus *trans*, we first considered the numbers of genes with only effects in *cis* (‘cis’ class), with only effects in *trans* (‘trans’), with both *cis* and *trans* (‘compensating’, ‘compensatory’, ‘overcompensating’, and ‘enhancing’), and with no change (‘conserved’). The distribution of these numbers across the 2x2 table was assessed for significance using a Fisher’s Exact Test (FET)/hypergeometric test. We also repeated this test excluding enhancing genes from the category of genes with both *cis* and *trans* effects (both were highly significant, all *p* < 2 x 10^-112^ and all *p* < 4 x 10^-102^, respectively). To assess the enrichment of opposing effects within the set of genes that showed both *cis* and *trans* regulation, we used a binomial test (always 87-92% of genes with *cis* and *trans* effects had those effects oppose each other across strains). To assess what proportion of genes called *cis-trans* opposing would have to be wrongly classified to result in the FET being not significant (after Bonferroni-correction), we sequentially removed 1-100% of these genes from this category in two ways, then re-computed the FET. First, to simulate DE being missed at a gene, we took genes away from the *cis-trans* opposing group and added them to the *cis* only group. Second, to simulate ASE being spuriously called at a gene, we took genes away from the *cis-trans* opposing group and added them to the conserved group. In each such simulation, we computed the proportion of all called DE genes simulated as being missed and all called ASE genes simulated as being spurious and report these numbers in the main text where the FET was no longer significant. This analysis script is in our code repository: *data_analysis_scripts/ compensationamounttesting.R*

#### Gene filtering

We performed all analyses including all nominally expressed genes, excluding genes overlapping hyperdivergent haplotypes or with aberrantly low or high DNA sequence coverage in the focal strain, and excluding all genes called hyperdivergent in any of 328 strains analyzed by CeNDR (Lee *et al*. 2021). Focal strain gene haplotype hyperdivergence was inferred when the gene region overlapped any hyperdivergent haplotype in this strain in the hyperdivergent haplotype BED file from the CeNDR 20210121 release (Lee *et al*. 2021). Genes were flagged as having aberrantly low or high DNA sequence coverage if they had <0.3 or >2.5-fold the median gene’s coverage in that strain, with coverage calculated across all exonic bases from CeNDR DNA sequence BAMs (20210121 release), as described previously (Bell *et al*. 2023). The list of genes hyperdivergent in any strain population wide was from Lee et al (Lee *et al*. 2021).

#### Gene set enrichment analyses

We used WormCat (Holdorf *et al*. 2020) to perform gene set enrichment analyses by writing a script extension to the WormCat R package (v2.0.1) that allowed us to provide a custom background gene set for enrichment tests. We performed the following tests with genes from each strain separately (formatted here as test gene set *vs* background gene set): DE genes *vs* all analyzed genes, ASE genes *vs* ASE-informative genes, compensatory genes *vs* ASE- informative genes, compensatory genes *vs* ASE genes, transgressive (overdominant + underdominant) genes *vs* all analyzed genes, overdominant genes *vs* all analyzed genes, underdominant genes *vs* all analyzed genes, DE genes that are ASE-informative *vs* ASE- informative genes, ASE-informative genes *vs* all analyzed genes, N2 dominant genes *vs* all analyzed genes, wild dominant genes *vs* all analyzed genes, *cis* genes that were not called additive inheritance mode *vs* ASE-informative genes, and *cis* genes that were not called additive inheritance mode *vs* ASE genes. The WormCat extension and analysis scripts are in our code repository: *data_analysis_scripts/wormcat_givebackgroundset.R* and *data_analysis_scripts/combinewormcatout_aseetc.R*.

#### Meta-strain results: combined comparisons across strains

We performed all analyses within each strain/strain pair, and we also combined strains’ results into one ‘meta-strain’ to be able to display and report one set of results when results across strains were largely consistent. For this meta-strain, genes were considered ASE-informative if they were ASE-informative in all seven strains and not ASE-informative if they were not informative for ASE in any strain; genes had to be informative in all strains or not informative in any strain to be compared in informative-vs-not analyses. To compare ASE vs. not, DE vs. not, and regulatory pattern, genes informative in all strains were included for each strain: each gene is present on each plot seven times, in the category of its classification for each strain. For example, one gene might be called ASE in three strains and not ASE in four strains and would be represented by three points in the ASE group and four points in the non-ASE group. In some cases, other characteristics of the gene (such as essentiality, see below) was the same across strains and therefore represented identically seven times while in others (such as expression level, see below) both the ASE characterization and the other characteristic are different in each strain.

#### Comparing to regulatory divergence between *C. nigoni* and *C. briggsae*

To determine how our estimates of regulatory divergence within *C. elegans* compare to the inter-species regulatory differences between nematode species *C. nigoni* and *C. briggsae,* we obtained the number of regulatory-diverged genes from the ASE analyses in Sanchez-Ramirez *et al*. (2021). In their df.cis_trans.inherit.merge.csv (https://github.com/santiagosnchez/competitive_mapping_workflow/raw/refs/heads/master/analyses/tables/cis_trans_and_expression_inheritance/df.cis_trans.inherit.merge.csv) dataset, 10,502 genes had assigned regulatory patterns, and 8653 (82.4%) of these had diverged regulation (*cis, trans*, or compensatory; ambiguous excluded). These species differ at 20.7% of synonymous sites (Thomas *et al*. 2015), equivalent to a strain differing from N2 at 20,756,530 bp. We used these numbers allowed and our within-species *C. elegans* linear model to predict the proportion of regulatory-diverged genes between *C. nigoni* and *C. briggsae* and compare that to the observed proportion of regulatory-diverged genes and to genomic divergence.

#### Genome, population genetic, and gene essentiality metrics

Genes were assigned to chromosome region bins (centers, arms, tips) based on which region from Rockman and Kruglyak (2009) contained the gene’s midpoint. Nucleotide diversity statistics population-wide pairwise segregating sites (π) and among-parental-pair proportion segregating sites *p* were calculated from the 20210121 hard-filter CeNDR VCF from biallelic SNVs only using PopGenome (v2.7.5) (Pfeifer *et al*. 2014). Nucleotide diversity (π) and Tajima’s D were also obtained from Lee *et al*. (2021), with their per-kb π per site converted to per-gene π per site by taking the median (missing data excluded) of all 1kb windows overlapping the gene +/- 500 bp. Tajima’s D, Fay & Wu’s H, and F_ST_ in non-Hawaiian and Hawaiian sub-populations were obtained from Ma *et al*. (2021). When we had multiple sources for the same statistic, we tested all of them, finding that results were generally consistent across statistic source when they were internally consistent across strains and gene sets; we use π from Lee *et al*. (2021) in the figures in this study. Whether the gene fell in a haplotype with a selective sweep in N2 was inferred from the swept haplotype data from Lee *et al*. (2021). To assign genes as essential or not, we downloaded gene annotations including “RNAi Phenotype Observed” and “Allele Phenotype Observed” for all genes in the *C. elegans* genome from WormBase using SimpleMine (Sternberg *et al*. 2024). Genes with lethality or sterility phenotypes from RNAi or alleles were considered essential (specifically, we searched for “lethal” and “steril” in the “RNAi Phenotype Observed” and “Allele Phenotype Observed” columns). Scripts used for these analyses are in our code repository:

*data_generation_scripts/nucdivcendr_geneswindows_allandasestrains.R,*

*data_analysis_scripts/chrlocenrichment_asederpim.R,*

*data_analysis_scripts/aseetc_vs_general.R*

#### Expression level analyses

For comparing gene categories to the expression level of each gene, we used the average normalized expression level from the six relevant parents in each cross. Specifically, kallisto quantification estimates to strain-specific transcriptomes were length and library size normalized followed by variance-stabilizing transformation (all via DESeq2), then averaged across the appropriate samples.

For analyses of human gene expression variability *vs* human gene expression level, we used the S4 dataset from Wolf *et al*. (2023), which comprises ranks of gene variation and expression level derived from principal components analysis of across-57-study correlation in gene expression variation and (separately) mean gene expression. Prior to this cross-study variance and level ranking, the authors corrected for the mean-variance relationship of gene expression within each study. We performed correlation tests on the input data as well as assigned genes to deciles of gene expression variability (1313 or 1314 genes per decile, 13139 genes in dataset), testing these deciles for differences in central tendency of gene expression level via ANOVA.

For analyses of *Drosophila melanogaster* gene regulatory pattern and variation vs expression level, we used data from Glaser-Schmitt *et al*. (2024). We obtained ASE, DE, tests of their difference, and regulatory pattern assignments for midgut and hindgut from Data S3 and Data S4, and raw gene counts from the related GEO repository (GSE263264). To generate normalized expression values to use in our analyses, we determined the length and GC content of each transcript in the *D. melanogaster* genome from the transcript FASTA from FlyBase (release FB2024_06) (Ozturk-Colak *et al*. 2024), and used the median across transcripts within a gene for the gene’s value. We corrected count data from all samples from the parent strains for length bias (highly relevant) and GC bias (not very relevant in *D. melanogaster*) using R package cqn (v1.48) (Hansen *et al*. 2012), then normalized to correct for library size and variance using DESeq2’s *vst* and *normTransform* functions. The point estimate for each gene’s expression was the mean across all parents’ normalized values in the specific cross; we did all comparisons with both *vst* and *normTransform* normalized data, which yielded qualitatively similar results. Downstream, we used *normTransform* data as it removed more of this data’s mean-variance correlation than did *vst*. To assign genes to regulatory patterns, we used Glaser-Schmitt *et al*. (2024)’s ASE and DE log_2_ fold changes and their Chochran-Mantel-Haenszel test p-values comparing the parental and F1 allelic ratios, then followed our regulatory pattern assignment criteria without requiring a log_2_ fold change threshold, as the original authors did not. Our regulatory pattern assignments were concordant with Glaser-Schmitt *et al*. (2024)’s except in cases where they hadn’t taken directionality of effects into account and where we called genes as conserved if they did not have ASE or DE as called by DESeq2, even if they had a significant CMH based on underlying allelic ratios. Because patterns were consistent across strains, we plotted all strains’ results together in. We focused our interpretations on genes with changed expression regulation as determined by their ASE and DE estimates and comparisons between them, as the threshold for ASE informative genes was different enough between the studies to preclude comparing genes without called expression changes.

Scripts used in these analyses are in our code repository:

*data_analysis_scripts/aseetc_vs_general.R, data_analysis_scripts/*

*wolf2023humexpanalyses.R, data_analysis_scripts/*

*glaserschmitt_drosase_meanexprvsaseetc.R*

#### General software tools used for analyses and figures

Tools used for specific analytical purposes are described in the relevant sections; here, we share tools used for general data processing and figure creation.

Analysis scripts were largely written in R (v4.3.2) (R Core Team 2023), with a few written in Python (v3.7) (www.python.org). Workflow scripts were written and run using Nextflow (v22.10.7) (www.nextflow.io). Compute-intensive analyses and workflows were run via the Partnership for an Advanced Computing Environment (PACE), the high-performance computing environment at the Georgia Institute of Technology.

General data wrangling R packages used included data.table (v1.14.99) (Dowle and Srinivasan 2022), argparser (v0.7.1) (Shih 2021), and formattable (v0.2.1) (Ren and Russell 2021). R packages used for data display and figure creation included ggplot2 (v3.5.1) (Wickham 2016), cowplot (v1.1.2) (Wilke 2020), eulerr (v7.0.2) (Larsson and Gustafsson 2018), ggforce (v0.4.1) (Pedersen 2022), ggVennDiagram (v1.2.3) (Gao 2021), ggsignif (v0.6.4) (PedersenAhlmann-Eltze and Patil 2021), and ggpmisc (v0.5.6) (Aphalo 2024). Color schemes were developed using RColorBrewer (v1.1-3) (Neuwirth 2022) and Paul Tol’s color palettes (https://personal.sron.nl/~pault/).

## Results

### An experiment to reveal extent and mode of gene expression variation in *C. elegans*

To interrogate intraspecific gene expression variation in *C. elegans*, we captured expression differences among the reference strain N2 and seven wild strains. We estimated pairwise differential expression (DE) between each wild strain and N2, as well as allele-specific expression (ASE) in the F1 offspring of each strain crossed to N2 (**Figure 1A, Table S1**). ASE analyses are uniquely sensitive at identifying *cis* regulatory changes (Cowles *et al*. 2002; Yan *et al*. 2002; Wittkopp *et al*. 2004), and analyzed in conjunction with DE of parental strains, they can reveal the regulatory pattern and inheritance mode of gene expression across the genome (**Figure 1B**). The seven wild strains were chosen to represent a range of nucleotide divergence from N2 and spanned the species’ genetic diversity: EG4348; DL238; CB4856 (‘Hawaii’); ECA722; QX1211; and ECA701 and XZ1516, two extremely diverged strains (**Figure 1C**).

**Figure 1.**
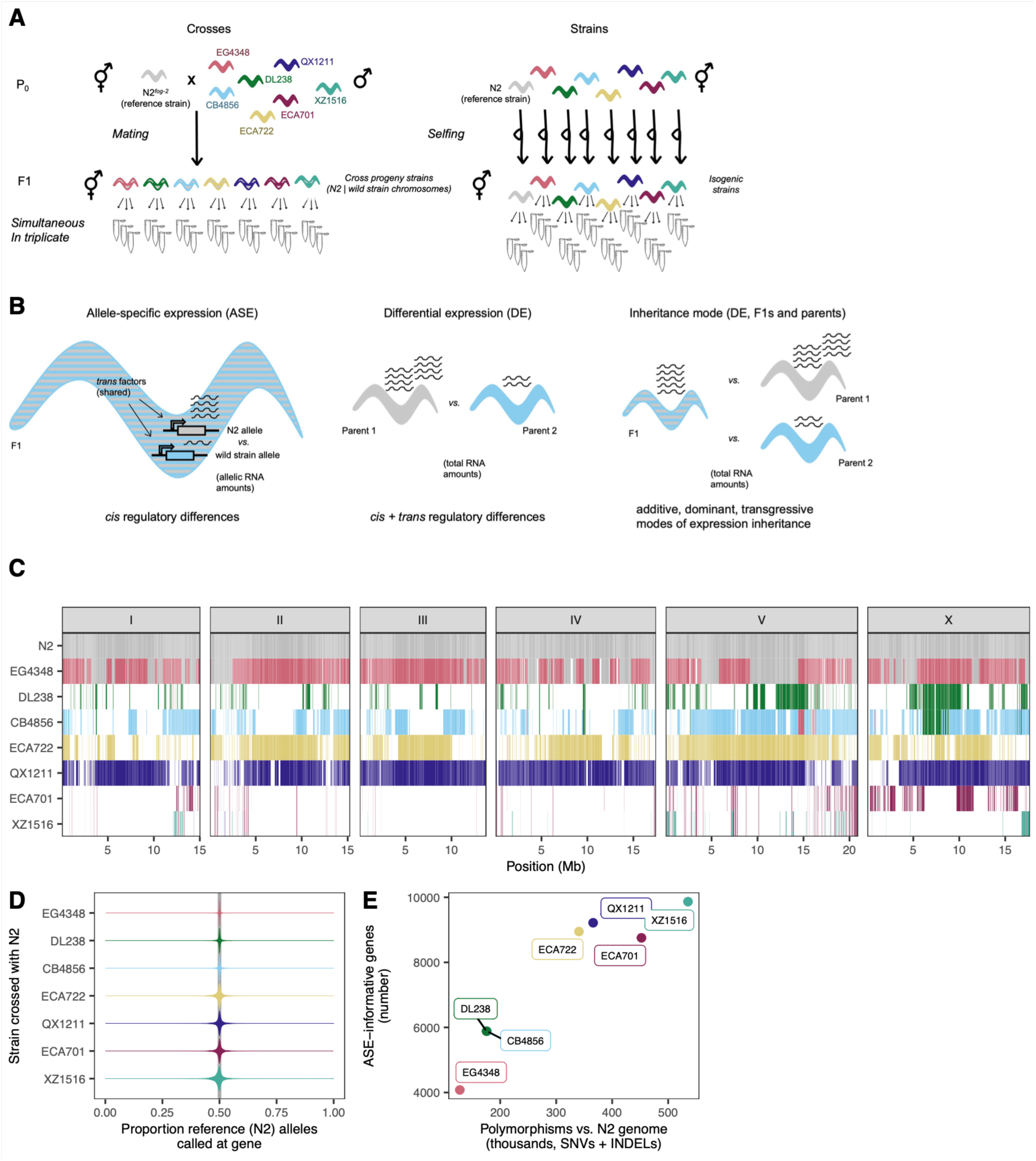
Interrogating gene expression variation in wild *C. elegans*. **A.** Experimental regime. **B.** The three expression level comparisons from this experiment. *Left*, allele-specific expression (ASE) is estimated from per-allele, allele-specific read quantification within each set of F1s. *Center,* comparison of total RNA amounts between parental strains yields differential expression (DE) estimates. Comparisons of ASE and DE enable determination of regulatory pattern of expression differences. *Right,* comparison of total RNA amounts between the F1 and its parents enables inference of inheritance mode of each gene’s expression. **C.** Genetic similarity of the strains in this study. White indicates haplotypes containing variants private to this strain in the entire population, whereas colors show haplotypes shared with at least one other strain in the population. Specific color denotes the first strain in this study in which the given haplotype was observed; the same color shows that haplotype as identical-by-descent with at least one other strain in the entire population (haplotype identity-by-descent data from Lee *et al*. (2021)). **D.** Proportion reference alleles in each ASE-informative gene’s RNA seq. (See Table S2 for all gene *n*s.) **E.** The relationship between number of ASE-informative genes (see main text) to the genome divergence between the wild parental strain and reference genome N2.

To maximize power and limit confounding effects, we conducted the experiment in one batch, generating young adult selfed offspring of the parental strains simultaneously with their cross offspring with N2 (**Figure 1A**, Methods). Replicate RNA-seq samples clustered in gene expression space, indicating true differences between strains and generations (principal components analysis, **Figure S1**). To analyze these gene expression data for signatures of DE and ASE, we developed a framework that 1) minimized reference bias, wherein sequence reads from the reference genome have higher rates of alignment than reads from the non-reference genome (Degner *et al*. 2009), 2) equivalently handled strains and genomes with varying levels of difference from each other while minimizing differences in power among strains as much as possible (**Note S1**), and 3) generated comparable estimates of expression differences among strains (DE) and between alleles within the F1s (ASE), enabling direct comparison (Methods). Although the wild strains exhibit a substantial span in their genetic differentiation from the reference, we observed no reference bias; the proportion of reference alleles called per gene was tightly centered around 50% for all strains (**Figure 1D**). To estimate DE among strains, we analyzed 18,647 genes with nominal expression (those with 10 or more total reads observed across samples). To detect ASE within the F1 hybrids, transcripts must carry genomic variant(s) that discriminate between the parental genotypes, so not all expressed genes permit ASE analysis. The genes informative for ASE comprised 22-53% of all nominally expressed genes; the proportion scales with genetic difference from N2 (**Figure 1E**). In this manuscript, we refer to these as “ASE-informative” genes.

Here, we present the insights derived from these gene expression data for all *C. elegans* genes, including those in hyperdivergent haplotypes (Lee *et al*. 2021; Moya *et al*. 2025), as global trends persisted across different gene inclusion criteria (Discussion).

### Modes of gene expression inheritance include transgressive variation

Of all analyzed genes, 26% exhibited differential expression within at least one cross, either between the parental strains or between generations (**Figure 2**). To evaluate how these expression changes were inherited, we compared, for each gene, the expression of the F1 offspring to each of its parents (McManus *et al*. 2010): we classified genes for which the F1 exhibits the same expression as one parent but different expression from the other as ‘N2 dominant’ or ‘alt dominant’ (wild strain dominant); genes with expression intermediate to the parents as ‘additive’; and genes with expression significantly higher or lower in the F1 than in both parents as transgressive, either ‘overdominant’ or ‘underdominant’ (**Figure 2A; Figure S2;** Methods). Each cross exhibited genes with each inheritance classification (**Figure 2B**), including overdominant and underdominant transgressive expression. Across strains, transgressive genes comprised 0.2-0.6% of all genes and 0.9-7.2% of genes with unambiguous expression changes. These transgressive genes are of outsized interest: expression values beyond the range of the parents may contribute to hybrid dysfunction and eventual speciation, and may help to explain outbreeding depression in *C. elegans* (Renaut *et al*. 2009; Gomes and Civetta 2015; Sanchez-Ramirez *et al*. 2021).

**Figure 2.**
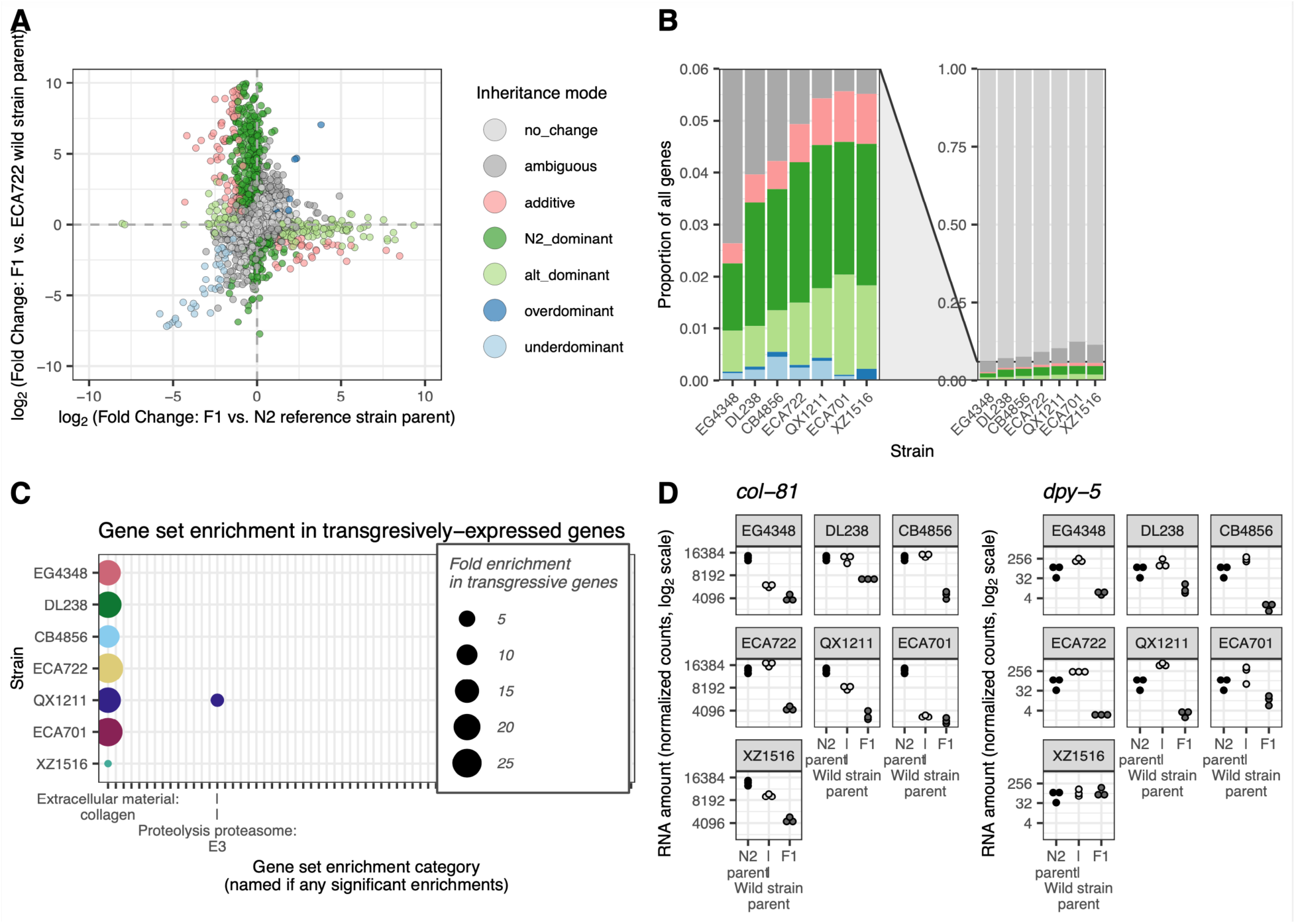
Gene expression inheritance modes. **A.** For each cross, the inheritance mode of each gene was inferred by comparing differential expression (DE) between the F1 and their N2 parent (x axis) and DE between the F1 and their wild strain parent (y axis) (McManus *et al*. 2010). One point per analyzed gene, excluding 20 exceeding the axis limits. *Figure S2 shows this classification for all strains.* **B.** Colors show the proportion of genes with each inheritance mode across strains (colors map to inheritance mode patterns as in A). Right: all genes; left: zoomed on genes with expression differences. *Table S2 has numbers (rather than proportions) of genes in each category in each strain cross*. **C.** Gene-set enrichment analysis results (Holdorf *et al*. 2020) for transgressively inherited underdominant genes, in which the F1 expression is lower than either parent, vs. all analyzed genes. X axis ticks indicate all gene categories analyzed in this comparison; only significant enrichments are labeled (Bonferroni-adjusted *p* < 0.05). *Figure S3 shows gene set enrichment analysis results for all analyzed inheritance mode gene sets.* **D.** An example of two collagen genes with underdominant expression in multiple strains. N2 parental gene expression is the same in each sub-plot (the same three N2 samples serve as the N2 parent for all strains). *n* = 45. Web app wildworm.biosci.gatech.edu/ase shows these plots and further information for any queried gene.

To determine whether expression of functionally related genes tends to be inherited the same way within and across strains, we performed gene set enrichment analyses (**Figure S3**) (Holdorf *et al*. 2020). Genes with transgressive expression were heavily and consistently enriched for collagen genes relative to all other categories (**Figure 2C**). Yet, the pattern of expression varied by gene and by strain. Some collagen genes were lower expressed in the F1 than in either parent in all strains, with either idiosyncratic intermediate expression in the wild parent (*e.g., col-81,* WBGene00000657) or consistently high expression in both parents (*e.g., dpy-5,* WBGene00001067) (**Figure 2D**). Strain XZ1516 often showed unique patterns, suggesting its collagen network may be strain-specifically regulated. At least some of the expression variation in collagen genes likely originates with the N2 genotype, which participated in each cross; N2 carries a derived mutation that modifies the phenotypic penetrance of cuticle mutations commonly used as markers in lab work (Noble *et al*. 2020). However, the differences by gene and expression patterns across strains suggest that collagen genes may be especially evolutionarily labile within this species, consistent with earlier work suggesting their rapid evolution and expression differentiation (Cutter and Ward 2005; Denver *et al*. 2005). Collagen genes interact in complex networks to form the worm cuticle (Higgins and Hirsh 1977; Cox *et al*. 1980; Kramer 1994; McMahon *et al*. 2003), and pathway architecture, including redundancies, may facilitate functional diversification across strains.

### Regulatory patterns reveal extensive *cis-trans* compensation within gene expression variation

We next sought to elucidate how expression differences are regulated in *cis* versus *trans*. For each ASE-informative gene, we compared the allele-specific difference in expression, which must occur in *cis*, to the expression difference between the parents, which may arise from regulation in *cis*, *trans*, or both (McManus *et al*. 2010). We classified genes with similar magnitude ASE and DE as regulated largely or solely in *cis* (‘cis’ category); genes with DE but no ASE as ‘trans’; and genes with DE exceeding ASE as ‘enhancing’. Genes with ASE but no or lower-magnitude DE reflect *cis* differences that are compensated by opposite-effect regulation in *trans*, producing either an incomplete offset of the *cis* effect (‘compensating’), a complete offset (‘compensatory’), or an opposite effect exceeding the change in *cis* (‘overcompensating’) (**Figure 3A; Figure S4;** Methods). This classification regime operates equivalently across strains, enabling inter-strain comparisons. Importantly, this method avoids a common pitfall wherein the influences of *cis* and *trans* effects are artifactually negatively correlated (**Note S2**; Fraser 2019; Zhang and Emerson 2019). Such correlation can inflate inferences of compensation because any false positive estimates of *cis* effects, leading to spurious calls of ASE, result in equal-but-opposite *trans* false positives to explain the absence of observed DE (**Note S2**; Fraser 2019; Zhang and Emerson 2019).

**Figure 3.**
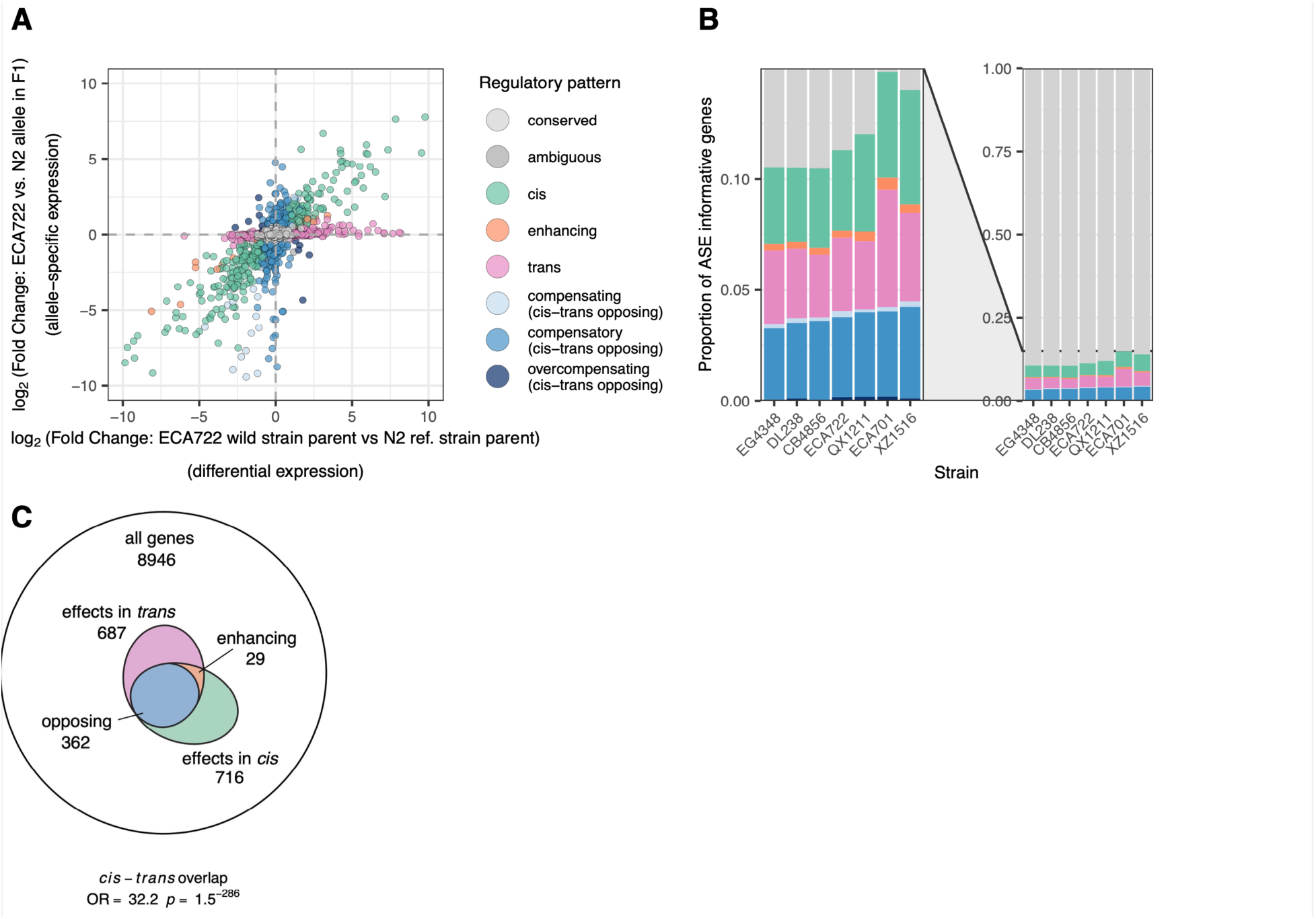
Gene expression regulatory patterns. **A.** The regulatory mode for each gene was inferred by comparing differential expression (DE) between the two parental strains (x axis) with allele-specific expression (ASE) between the two alleles in the F1 (y axis) (McManus *et al*. 2010). One point per ASE informative gene, excluding 10 exceeding the axis limits. *Figure S4 shows this classification for all strains.* **B.** Colors show the proportion of genes with each regulatory pattern in each strain cross (colors map to regulatory patterns as in A). Right: all genes; left: zoomed on genes with expression differences. *Table S2 shows numbers (rather than proportions) of genes in each category in each strain.* **C.** Relative to all genes, the subset of genes with regulatory differences in *cis* and in *trans* (in isolation or in combination), shown for strain ECA722; many more genes have both *cis* and *trans* effects than would be expected by chance if the effects are independent (hypergeometric/Fisher’s Exact Test results in panel). Numbers show total number of genes in category inclusive of category-specific and overlapping genes. *Figure S6 shows this result for all strains*.

Across strain pairs, 9-15% of the ASE-informative genes exhibited expression differences in *cis*, *trans*, or a combination; roughly similar numbers of genes were regulated primarily in *cis*, primarily in *trans*, or both in opposite directions indicating compensatory regulation (**Figure 3B**). As expected, the genes with no observed expression differences in the inheritance mode analysis were likewise classified as unchanged (‘conserved’) or as compensatory in this analysis (**Figure S5**). Genes with expression differences driven solely in *trans* were much more likely to be inherited as dominant than as additive, and likewise genes with expression differences driven solely in *cis* were more likely than *trans* genes to be inherited additively. This trend makes sense, as a *trans* factor inherited from one parent can target alleles from both while a change in *cis* affects only the local allele (Lemos *et al*. 2008). However, many *cis* regulated genes were also inherited dominantly, as has been observed in other nematode species (Sanchez-Ramirez *et al*. 2021). While this result may reflect real biological mechanisms (Sanchez-Ramirez *et al*. 2021), we note that estimates of dominance may be inflated since the statistical threshold for additivity, which requires the intermediate F1 expression level to be distinct from both parents, is harder to achieve than that for dominance, which requires distinction from only one parent.

The substantial fraction of genes exhibiting compensatory regulation (**Figure 3B**) reflects the ubiquity of opposing *cis* and *trans* effects at the same individual genes. For example, of genes with detected ASE, *i.e.*, evidence of gene regulation in *cis*, 44-51% were inferred to be partially or completely attenuated in *trans*, resulting in no or reduced differential expression at the organismal level (**Figure 3C, Figure S6**). This is a substantially higher overlap than would be expected by chance if the probabilities of *cis* and *trans* effects were independent (FET, all *p* < 2 x 10^-112^, **Figure S6**). Moreover, of genes regulated in both *cis* and *trans*, opposing effects (the compensatory classes) dominated, comprising 88-94% of these genes. This significant excess of genes with *cis*-*trans* overlapping effects is unlikely to be artifactual: for this enrichment of compensatory effects not to be significant, 90% of these calls would need to be in error, meaning at least 40% of our ASE calls would have to be spurious (false positives) or 35% of DE calls would have had to have been missed (false negatives) (see also **Note S2**). Such pervasive opposition between regulatory effects may reflect a history of stabilizing selection on gene expression levels (Discussion).

We next considered whether the expression differences we observed, and their modes of regulation, were common or idiosyncratic across strains. As all crosses shared N2 as a parent, expression differences arising from derived changes in N2 are likely to be shared; this is evidenced by the shared expression pattern at *fog-2* (WBGene00001482), which exhibited allele-specific expression in each cross (we deleted this spermatogenesis gene from the N2 worms used to create the F1s, to facilitate obligate outcrossing, though we used wild-type N2 for sequencing this ‘parental’ strain). Strain pairs showed consistency in expression variation with respect to which genes exhibited any differences at all: 1798 (85%) of shared informative genes had the same regulatory pattern at 6 or more strains, and 1480 (70%) at all 7 strains (**Table S3**); most of these genes shared conserved expression. However, of genes with expression differences, 82 exhibited *cis* dominant regulation in at least one strain and *trans* dominant regulation in at least another strain, and 109 genes exhibiedt uncompensated *cis* driven expression changes in at least one strain that were compensated in at least one other strain (**Table S3**). While apparent differences among strains may be inflated by the statistical challenge of observing a given gene as significant in multiple crosses, the overall trend is that genes with expression differences were strain specific: of genes that were ASE-informative in all strains, a preponderance (51.9%, 275 of 530) of those exhibiting ASE did so in only a single strain (**Figure S7, Table S3**). Genes with specific regulatory patterns tended not to be obviously associated with common biological categories across strains (**Figure S8**).

### Expression variation increases with genomic divergence

*C. elegans* strains persist predominantly as selfing lineages, resulting in the accumulation of genetic changes and a spectrum of genomic differentiation between more closely or more distantly related strains (Barriere and Felix 2005b; Barriere and Felix 2005a). We leveraged this aspect of *C. elegans* biology to assess the relationship between genomic differentiation and gene expression variation, asking whether the proportion of genes with expression differences changes with genomic differentiation. Overall, yes: for each strain, the proportion of genes with differences in expression scaled positively with genetic distance from N2 (**Figure 4**).

**Figure 4.**
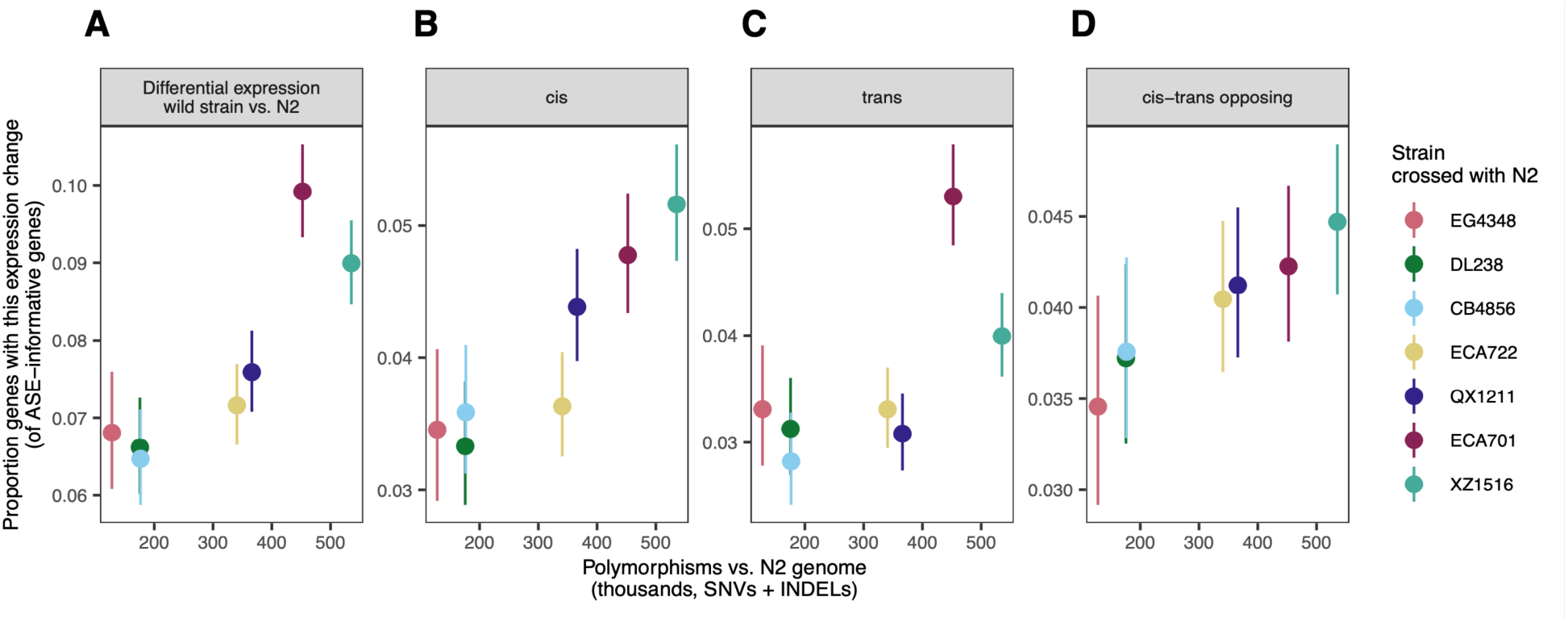
Increasing expression variation with genomic divergence. For each wild strain, the proportion of genes exhibiting a given regulatory difference is plotted against the genetic distance from reference strain N2. **A,** Differential expression between the wild strain and N2 at ASE informative genes; **B-D,** proportion of genes with specific regulatory patterns in each strain cross). *Cis-trans* opposing genes comprise compensating, compensatory, and overcompensating gene categories from Figure 3A,B. All non-conserved expression regulatory patterns combined: correlation ρ = 0.89, *p* = 0.01. In all panels, error bars denote 95% binomial confidence intervals of proportion and y-axis bounds are specific to the data shown. *Figure S9 shows proportion of each individual regulatory pattern category vs. genetic distance from N2; Figure S10 shows proportion of each individual inheritance mode vs. genetic distance from N2.* See Table S2 for all gene *n*s.

This pattern persisted across all measures of expression variation: for genes exhibiting DE between strains (**Figure 4A**), for genes by regulatory class (**Figure 4B-D, Figure S9**), and for genes by inheritance mode (**Figure S10**), with weaker trends for the infrequently observed gene categories with less estimate precision. Considering all genes with evidence of differential regulation (all those not called conserved), we estimate that increasing the number of genetic variants by 100 thousand increases the proportion of variable expression genes by one percentage point (1%) (linear regression per 1000 variants: β = 1.2 x 10^-4^, *p* = 0.004 p). This trend is not explained by the increased number of ASE-informative genes in more highly differentiated strains, as the estimates are specific to the ASE-informative genes for each strain (though see **Note S1**).

Given that *C. elegans* live as independent lineages and show significant outbreeding depression (Dolgin *et al*. 2007), we had hypothesized that gene expression variation might plateau between the most diverged strains, as observed between species. For example, the nematode species *C. briggsae* and *C. nigoni* exhibit regulatory divergence at 82% of ASE-analyzed genes (Sanchez-Ramirez *et al*. 2021), which is 13.5-fold more expression differences than our most-diverged *C. elegans* strain pair even as these sister species are 39-fold more diverged at the genome level (Thomas *et al*. 2015). However, here, gene expression differences scale with genomic differentiation and do not show any softening of the trend among the most diverged strains, as we might expect between incipient or distinct species.

### Location, nucleotide diversity, and essentiality define genes with expression differences

To consider processes that may have shaped gene expression variation, we interrogated gene sets with different regulatory patterns for association with genomic location, nucleotide diversity metrics, and gene essentiality.

The *C. elegans* genome harbors extensive evidence of its recombination history, with more recombination in the chromosome arms and less in chromosome centers (Rockman and Kruglyak 2009): gene density tends to be higher in the centers while nucleotide diversity is higher on arms (Rockman and Kruglyak 2009; Andersen *et al*. 2012). ASE-informative genes must have coding sequence polymorphisms; commensurately, they are enriched in chromosome arms and exhibit higher nucleotide diversity across all strains (**Figure 5A-B; Figure S11-13**). However, even accounting for this background enrichment, genes with expression differences (in *cis* or *trans*) were more likely to reside on chromosome arms than on centers (**Figure 5A, Figure S11**). These trends held for all seven strains and reinforce prior observations that genes variably expressed across wild *C. elegans* strains are more likely to reside in arms, as shown for differential expression (Denver *et al*. 2005) and as mapped as eQTLs by linkage (Rockman *et al*. 2010) or by association (Zhang *et al*. 2022). Relatedly, genes with expression differences showed an excess of polymorphism—beyond that which makes them ASE-informative—and were associated with both elevated genetic differences between the two parents (**Figure S12**) and elevated nucleotide diversity across the species (**Figure 5C, Figure S13**). This trend parallels recent findings in humans that genes with higher variation in expression harbor more genetic polymorphism (Wolf *et al*. 2023). This trend likely reflects differences in historical selection, including the possibility of selection relaxation (Discussion); consistent with this, expression-stabilized genes exhibiting compensatory regulation with opposing *cis* and *trans* effects had lower nucleotide diversity, and tended to be less enriched in chromosome arms, than non-compensated genes, but not to the extent shown by genes with completely conserved expression (**Figure 5A-C**).

**Figure 5.**
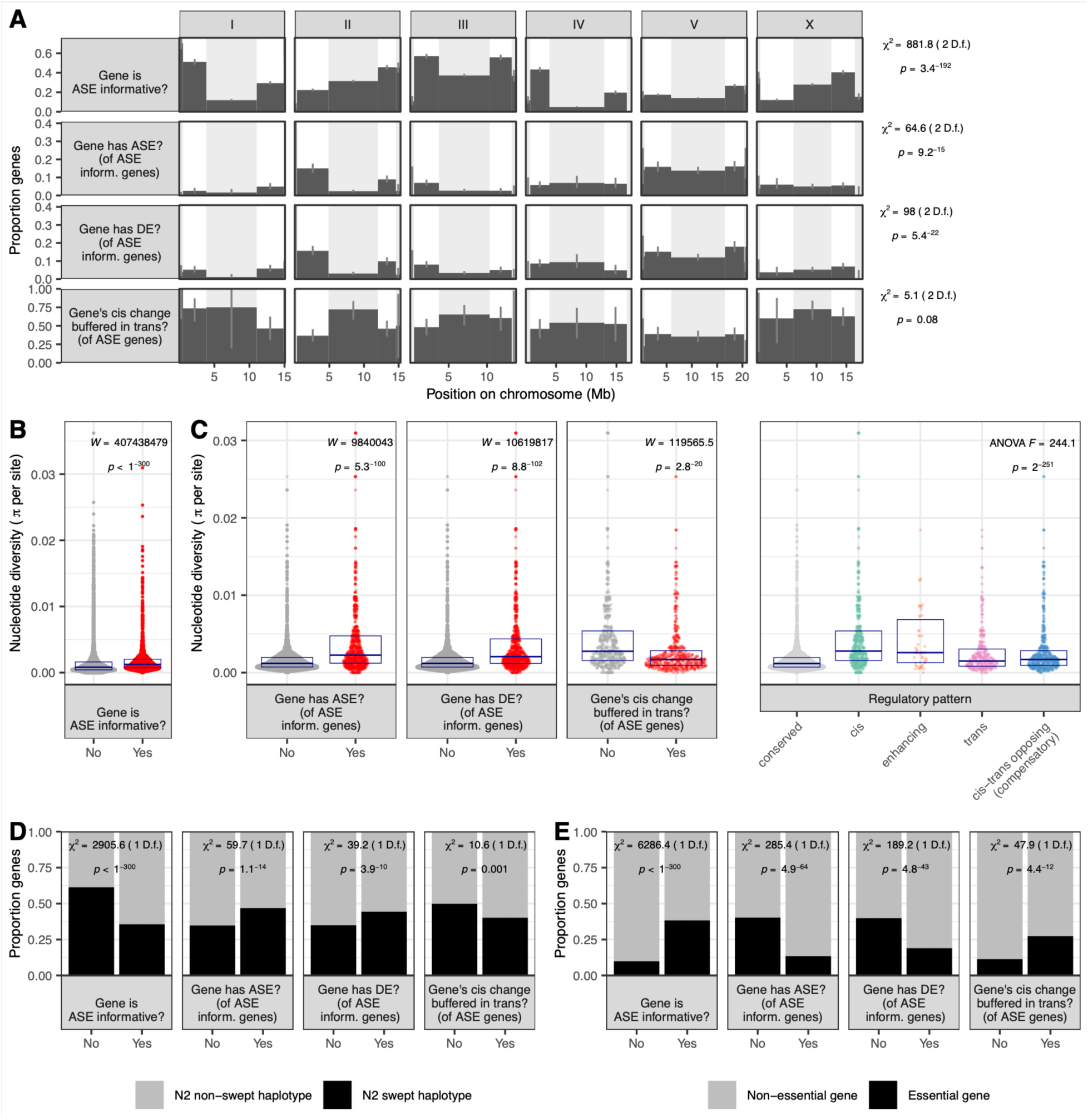
Location, nucleotide diversity, haplotype, and essentiality differentiate expression diverged genes. Results shown here are for all strains combined (Methods). See Table S2 for all gene *n*s. **A.** Proportion of genes in each region of the chromosome (tip, arm, and center, denoted by alternating white and gray background) that have the described attribute. *Figure S11 shows similar data for all strains individually.* **B-C.** Average per-site nucleotide diversity, estimated species-wide from 300+ wild *C. elegans* strains, is shown for genes with various expression patterns. Each point represents one gene and points fill a violin plot; boxes denote median +/- interquartile range. In **C**. **(right)**, Tukey’s HSD on annotated ANOVA cis > conserved (*p* = 9.8 x 10^-9^); enhancing > conserved (*p* = 9.8 x 10^-9^); trans > conserved (*p* = 9.8 x 10^-9^); cis-trans opposing > conserved (*p* = 9.8 x 10^-9^); cis > trans (*p* = 9.8 x 10^-9^), *cis* > cis-trans opposing (*p* = 9.8 x 10^-9^), enhancing > trans (*p* = 4.5 x 10^-5^), enhancing > cis-trans opposing (*p* = 0.0003) (all *p* values Bonferroni corrected; other comparisons non-significant). *Figure S12 shows pairwise, rather than population-wide, nucleotide diversity for all strains individually. Figure S13 shows same population-wide nucleotide diversity data for all strains individually.* **D.** For various expression characteristics of interest, bars show the proportion of genes in a region in which there is evidence of historical positive selection (selective sweep) in the N2 parent. *Figure S14 shows this breakdown for each strain individually.* **E.** As in **D**, but each bar shows the proportion of genes in that category that are predicted to be essential in *C. elegans*. *Figure S15 shows this breakdown for each strain individually*.

The *C. elegans* genome shows evidence of selective sweeps, in which haplotypes comprising large portions of individual chromosomes have risen in frequency across the population (Andersen *et al*. 2012; Lee *et al*. 2021). A footprint of strong historical selection, these sweeps dominate the genomes of non-Hawaiian isolates and may underlie adaptation associated with the colonization of new habitats (Zhang *et al*. 2021). We hypothesized that swept haplotypes are also associated with changes to gene expression. In our study, the non-Hawaiian strains N2 and EG4348 carry swept haplotypes over 65% and 37% of their genomes, respectively; the other Hawaiian strains harbor no swept haplotypes (Lee *et al*. 2021). Therefore, all our F1s share swept haplotypes inherited from N2, and only F1s derived from EG4348 carry additional swept haplotypes. Across strains, ASE-informative genes were less likely to reside in locations associated with N2 swept haplotypes (**Figure 5D, Figure S14**). However, genes with *cis* regulatory differences (ASE) and genes with expression differences (DE) were both more likely to reside in locations associated with sweeps in N2 (**Figure 5D, Figure S14**). These expression differences may have helped drive shifts in allele frequency and facilitated adaptation as *C. elegans* lineages colonized new habitats (Zhang *et al*. 2021), though the actual targets of selection within the swept haplotypes are unknown. Genes with *cis* regulatory differences compensated in *trans* tended to be less likely to be associated with swept haplotypes, but these trends were not always statistically significant across strains and gene sets (**Figure 5D**; **Figure S14**).

Next, we asked whether gene essentiality was associated with differences in expression. Essential genes, defined as those with an RNAi or allele phenotype leading to lethality or sterility (Sternberg *et al*. 2024), were significantly depleted among genes with any observed expression differences (ASE or DE), even as informative genes were enriched for essentiality (**Figure 5E**; **Figure S15**). These results reinforce earlier findings that essential genes are depleted among eQTL genes (Rockman *et al*. 2010; Zhang *et al*. 2022) and parallel observations from humans that genes with less expression variability tend to be less tolerant of loss of heterozygosity (Wolf *et al*. 2023). However, genes with opposing effects in *cis* and *trans* tended not to be depleted for essential genes (**Figure 5E; Figure S15**). Essential genes are therefore likelier to have expression differences compensated by another mechanism, stabilizing their expression.

### Genes with expression differences are less highly expressed

Finally, we examined the relationship between overall expression level and the tendency of genes to show expression differences. As higher expression enables the detection of significant differences, a positive association might arise as an artifact of the method; in fact, genes informative for ASE were higher expressed than those not ASE-informative (**Figure 6A, Figure S16**). Alternatively, a negative association between expression level and expression variation might reflect a phenomenon arising from biological function or evolutionary history.

**Figure 6.**
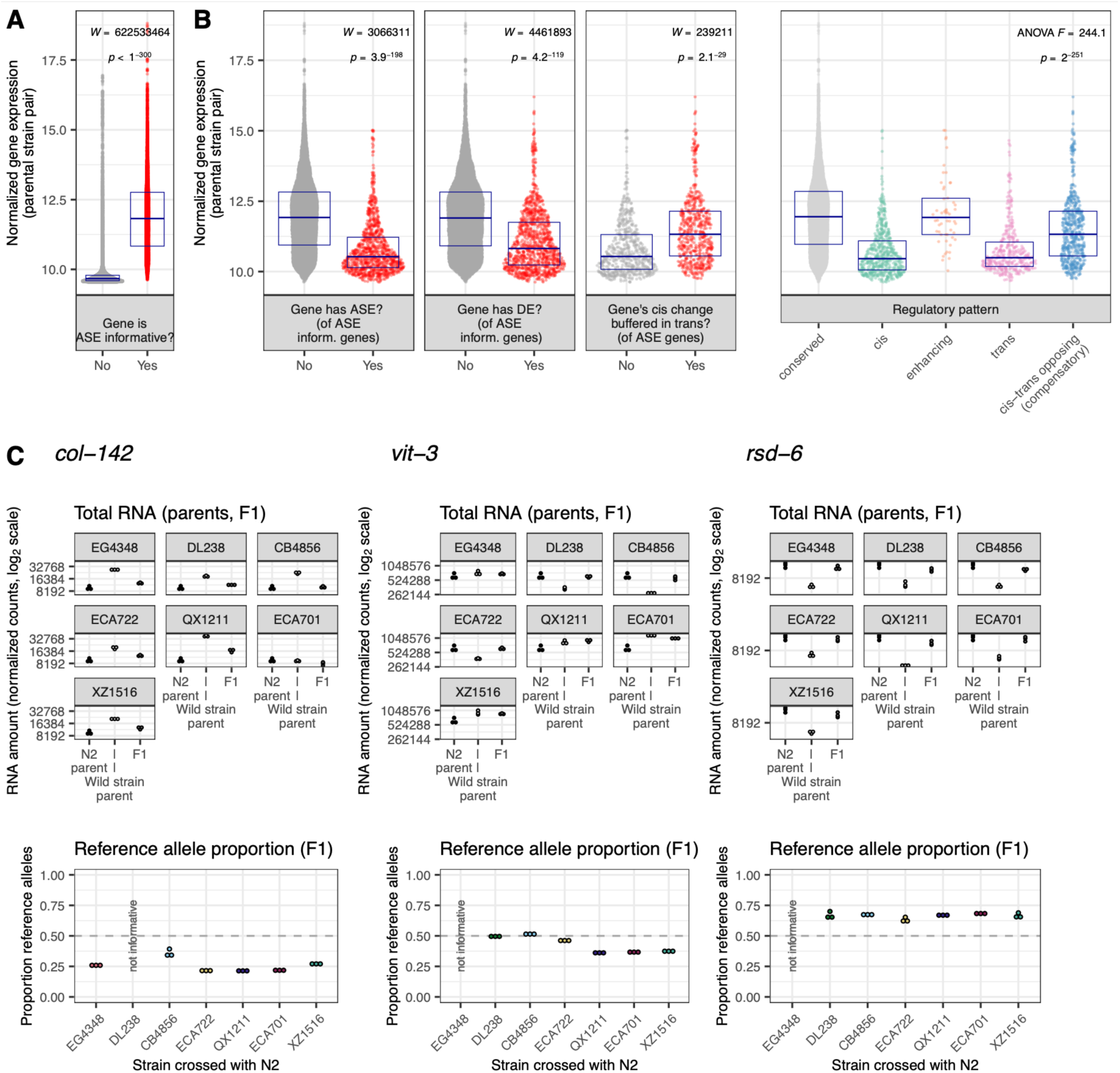
The relationship between expression level and expression variation. **A-B** Results shown are for all strains combined (Methods). Y axis denotes gene expression amount (length and library size normalized and variance stabilized, averaged across the two parental strains). Each point represents one gene and points inhabit a violin plot; boxes denote median +/- interquartile range. See Table S2 for all gene *n*s. In **B.** (**right**), ANOVA Tukey’s HSD conserved > cis (*p* = 9.6 x 10^-9^); conserved > trans (*p* = 9.6 x 10^-9^); conserved > cis-trans opposing (*p* = 9.6 x 10^-9^); cis > enhancing (*p* = 9.6 x 10^-9^), cis-trans opposing > *cis* (*p* = 9.6 x 10^-9^), *trans* > enhancing (*p* = 9.6 x 10^-9^), enhancing > cis- trans opposing (*p* = 0.014); (*p* = 9.6 x 10^-9^), cis-trans opposing > trans (*p* = 9.6 x 10^-9^), (all *p* values Bonferroni corrected; other comparisons non-significant). *Figure S16 shows expression vs. these various gene categories for all strains individually.* **C.** Three example *C. elegans* genes with top 10% expression levels that nonetheless exhibit DE caused by *cis* regulatory divergence. Top: total gene expression for each sample. N2 samples are the same across plots/crosses. Bottom: within-sample allelic proportion from allelic counts. *n* = 3 per strain per generation (45 total). Web app wildworm.biosci.gatech.edu/ase shows these plots and further information for any queried gene.

Here, genes with expression differences tended to exhibit lower average expression: of ASE- informative genes, those with detected ASE or DE were on average less expressed than those without (**Figure 6B, Figure S16**). Moreover, genes with no or reduced differential expression between strains because of *cis-trans* opposing effects exhibited intermediate expression levels, higher than uncompensated genes with expression differences but lower than conserved expression genes (**Figure 6B, Figure S16**). Because these genes with compensatory effects have higher expression than those with DE or uncompensated ASE, these calls are unlikely to be explained by missed DE calls or spurious ASE calls. To our knowledge, these are the first observations to explicitly demonstrate a relationship between gene expression level and gene expression variation, though genes with higher expression tend to evolve more slowly over interspecific timescales (Zhang and Yang 2015; Liao and Zhang 2006; Pal *et al*. 2001; Krylov *et al*. 2003). Thus, our results may reflect, at the intraspecific level within *C. elegans*, evidence of the constraint that has been hypothesized to govern the anticorrelation between expression and evolutionary rate (Zhang and Yang 2015) (Discussion). Because this pattern was clear in each strain, it is likely a general feature of *C. elegans* gene expression rather than a strain-specific idiosyncrasy (**Figure S16**).

To evaluate whether this relationship between gene expression level and variability extended beyond *C. elegans,* we examined expression data from humans and *Drosophila melanogaster* (**Figure S17**). For humans, we re-analyzed data from a meta-analysis of gene expression studies, comprising 57 studies with a median of 251 individuals included per study, which ultimately generating a robust across-study rank of mean expression and expression variance for each gene that encompassed variation driven by genotype and other sources (Wolf *et al*. 2023). In our analysis, more variably expressed human genes tended to be less expressed; the relationship is modest in magnitude but statistically significant (**Figure S17A**). For *D. melanogaster*, we re-analyzed and carefully normalized data from a recent allele-specific expression study of intestinal tissues from four strain crosses (Glaser-Schmitt *et al*. 2024). As we saw in *C. elegans*, *D. melanogaster* genes with compensatory *cis-trans* opposing effects as well as enhancing *cis-trans* effects had higher average expression than genes regulated only in *cis*, but *trans* effects were also elevated (**Figure S17B**). How much of these similarities and differences derive from study-specific analysis pipelines versus species-specific evolution remain open questions.

The observation that differentially expressed genes have lower expression on average provides a platform for identifying potentially important outliers: genes with very high expression that nonetheless have expression differences might be targets of strain-specific adaptive evolution. Of genes in the top 10% of gene expression, nine had *cis* regulated differential expression in one or more strains (**Table S4**). Anecdotally, these genes reflect dominant aspects of *C. elegans* biology: first, collagen genes *col-8* (WBGene00000597) and *col-142* (WBGene00000715, **Figure 6C**) are part of the extensive, epistatic network of genes coding for the collagen cuticle matrix. Second, vitellogenin genes *vit-3* (WBGene00006927, **Figure 6C**) and *vit-5* (WBGene00006929) code for extremely highly expressed yolk proteins that dominate young adult *C. elegans’* mRNA and protein generation (Perez and Lehner 2019) and whose gene products are even hypothesized to be used for offspring provisioning as a sort of ‘milk’ (Kern *et al*. 2021). Third, *rsd-6* (WBGene00004684, **Figure 6C**) and *deps-1* (WBGene00022034) are involved in the P granule and piRNA processing (Grishok 2013; Sternberg *et al*. 2024). Such small RNA pathways predominate worm biology and exhibit remarkable diversity in function and gene makeup across strains (Youngman and Claycomb 2014; Felix 2008; Chou *et al*. 2024). Although these genes exhibit similar high expression level and expression regulation, they are likely shaped by different evolutionary histories. For example, *rsd-6* is expressed at a lower level in all wild strains than in N2, suggesting an N2-specific mutation or function at this gene, while *vit-3* exhibits a diversification of expression levels across strains.

## Discussion

### Main findings

Our study of intraspecific variation in gene expression included the first allele-specific analysis in *C. elegans* and elucidates the gene regulatory architecture of this system. We observed substantial compensatory regulation, in which opposite effects in *cis* and *trans* at individual genes mitigate expression differences among strains. The most common gene expression pattern was one of no change and, while this majority gene set was well conserved among strains, genes with expression differences exhibited strain specificity and diversity in regulatory classification. We also report for the first time that expression-variable genes are lower expressed on average than genes without expression differences. Additionally, the number of expression-variable genes between strains increased with genomic distance. In addition to these broad trends, we also highlighted genes with outlier expression patterns, such as collagen genes heavily enriched for transgressive variation (**Figure 2D**) and expression-variable genes with extremely high average expression (**Figure 6C**).

The compensation of gene expression changes by *cis*-*trans* opposing interactions (**Figure 3**) has been observed across the tree of life, though with varying degrees of quantitative characterization and control for methodological artifacts (**Note S2**) (Goncalves *et al*. 2012; Coolon *et al*. 2014; Mack *et al*. 2016; Fear *et al*. 2016; Verta *et al*. 2016; Metzger *et al*. 2017; Fraser 2019; Zhang and Emerson 2019). While these interactions are often interpreted as evidence for stabilizing selection on expression level, recent work in yeast has demonstrated how an initial mutation, by skewing the mutational target space, may predispose later mutations to induce opposing, compensatory effects on gene expression (PedersenMcQueen *et al*. 2025). Under this regime, compensatory changes may accumulate neutrally, providing an alternative explanation for the significant excess of opposite effects relative to enhancing effects we observe in genes with changes in both *cis* and *trans*. Nevertheless, we found that genes exhibiting compensatory regulation were more likely to be essential than genes with uncompensated expression changes, suggesting stabilizing selection on expression. *C. elegans* may be especially prone to fitness-associated compensatory evolution due to extensive linkage across the genome arising from its predominantly selfing mode of reproduction (Barriere and Felix 2005b; Barriere and Felix 2005a; Rockman and Kruglyak 2009): fitness in *C. elegans* is mediated by opposite-effect, closely linked regions of the genome (Bernstein *et al*. 2019), and compensatory regulatory elements are closely linked in self-fertilizing spruce trees (*cis-trans*) (Verta *et al*. 2016) and yeast (*trans-trans*) (Metzger and Wittkopp 2019).

Our discovery that genes with expression differences tended to be expressed at lower levels (**Figure 6**) may have been overlooked previously given that most studies control for the positive correlation between mean and variance in RNA quantification, discouraging investigation into the larger phenomenon. Indeed, breaking this relationship is difficult, which limited our ability to assess this phenomenon in published data from other systems. However, we did also see evidence of this trend in humans and *D. melanogaster* (**Figure S17**). These sparse findings may relate to the more widely observed, and more thoroughly considered, phenomenon in which highly expressed genes evolve at a slower evolutionary rate (Krylov *et al*. 2003; Liao and Zhang 2006; Pal *et al*. 2001; Zhang and Yang 2015). This expression-rate anticorrelation is universal across the domains of life and has been explained by a number of mechanisms independent of selection on biological essentiality: avoidance of protein misfolding or mis-interaction, selection on mRNA folding, and cost of protein expression (Zhang and Yang 2015). Very few studies have explicitly investigated this phenomenon over micro-evolutionary timescales, but venom protein expression in pit vipers evolves more slowly, *i.e.*, is less variable, at higher levels (Margres *et al*. 2016), and higher-expressed genes in yeast exhibit lower nucleotide polymorphism due to purifying selection directly targeting amino acid identity (Marek and Tomala 2018). We posit that our observed anticorrelation of gene expression level and variability is likely a function of stabilizing selection on higher expressed genes in *C. elegans*, but the fitness-associated traits under selection may relate to the efficiency or efficacy of gene products rather than, or in addition to, ultimate biological function. We also note that the mutations that define the expression-rate anticorrelation must arise within populations, and exploration of the question within a tractable experimental system enables tests of longstanding hypotheses about the driving mechanisms (Biesiadecka *et al*. 2020). Overall, the expression level-variability anticorrelation invites many questions, including whether it occurs as a common feature of heritable expression variation in most systems; whether it scales to interspecific differences in expression; whether and how it translates to other molecular phenotypes, such as the expression of proteins; and the proximate and evolutionary forces that govern it.

We found that as genomic differentiation between the wild strains and N2 increased, the proportion of genes with expression differences also increased (**Figure 4**; see also **Note S1** for potential caveats). Such a trend is not necessarily to be expected: regulatory divergence has been observed to scale with genetic divergence among marine-freshwater ecotypes in sticklebacks (Verta and Jones 2019) but to plateau at high genetic divergence between yeast species (Metzger *et al*. 2017) and to not necessarily increase with divergence within and among *Drosophila* species, but to accelerate in particular crosses (Coolon *et al*. 2014). Though analyses of this relationship between genomic and regulatory divergence can shed light on the evolution of the genotype-phenotype map and the interplay between genetic variation, gene expression, and speciation (Mack and Nachman 2017; Orr 1995), it remains incompletely understood. Ultimately, this correlation must plateau at sufficient evolutionary distance, as genomic differences accumulate but not all genes show expression divergence; while 82% of genes exhibit regulatory divergence between sister species *C. briggsae* and *C. nigoni* (Sanchez-Ramirez *et al*. 2021), their genomic distance would need to be 2.9 times higher to scale with the trend we see within *C. elegans*. While it is difficult to directly compare ASE studies due to differences in experimental and analytical frameworks, we note similarities in the inheritance modes and regulatory patterns observed within *C. elegans* and between *C. briggsae* and *C. nigoni*, including the extent of compensatory regulation (Sanchez-Ramirez *et al*. 2021). The relationship between gene expression divergence and genomic divergence within *C. elegans* may offer an access point for deeper investigation within a highly tractable genetic system.

Consistent with earlier work testing the neutrality of *C. elegans* gene expression variation using mutation accumulation lines (Denver *et al*. 2005), our observations suggest that the majority of *C. elegans* genes may be under stabilizing selection for expression level. Genes with expression changes exhibit higher nucleotide variation, lower average expression, and a depletion of essentiality, potentially reflecting a history of relaxed selection relative to genes with stabilized expression. This result also comports with the observation that population structure estimated from gene expression data of 207 *C. elegans* wild strains is less differentiated than that estimated from nucleotide data, suggesting possible stabilizing selection on expression relative to nucleotide differentiation (Zhang *et al*. 2022), and with a recent analysis that inferred stabilizing selection at most *C. elegans* transcripts (PedersenInskeep and Groen 2025). However, in our results, genes with expression changes were also more likely to reside on chromosome arms than centers; this pattern may be explained in part by stronger stabilizing selection on expression in center-located genes but is also likely shaped by background selection, which acts independently of the expression phenotype of any individual gene (Rockman *et al*. 2010). Consequently, the role of stabilizing selection on expression phenotypes versus the indirect influence of background selection on reducing standing variation in chromosomal centers is not clear. Our observation that genes in chromosome centers show elevated compensation of expression differences by *cis-trans* opposing effects (**Figure 5A**) suggests that these may be direct targets of stabilizing selection.

We also observed evidence for adaptive evolution of gene expression variation. Namely, genes with expression differences were more likely to reside in locations at which the N2 haplotype experienced a selective sweep, which may include genes that facilitated adaptation during colonization of new habitats (Zhang *et al*. 2021). However, as in the case of background selection, such associations with genomic location do not distinguish between direct targets of selection and those indirectly targeted through linkage. Moreover, strain-specific changes in gene expression, even as they are associated with higher nucleotide diversity and other markers of relaxed selection, may instead reflect adaptive diversification; we cannot distinguish between these alternatives. For genes involved in environmental sensitivity or immune response, for example, the lower expression of expression-variable genes may be mediated by the absence of pathogens or other inducible factors in the lab environment. Thus, genes with expression differences yet high average expression may be useful candidates for identifying fitness-associated traits governed by expression variation.

In our study, each wild strain was crossed to the common reference strain N2, so N2-specific differences such as laboratory-derived adaptations would likely show up as common differences across the strain set. We observed only a small number of genes with common differences across all wild strains; instead, many genes had expression differences only in a single wild strain (**Figure S7, Table S3**). Genes in the worm cuticle network exhibited both shared and strain-specific trends. For example, most wild strains showed transgressive expression at the same collagen genes (**Figure 2C-D**), suggesting N2-specific differentiation. This result may relate to the derived mutation in *col-182* in N2, which increases the phenotypic penetrance of classical lab mutations affecting cuticle phenotype (such as *rol-1*) that are suppressed in the ancestral background (Noble *et al*. 2020). However, strain XZ1516 and its F1s exhibited distinct collagen gene expression phenotypes, suggesting divergent evolution in collagen or cuticle pathways along the XZ1516 lineage. The collagen gene network is especially large and complex with evidence of accelerated adaptation (Cox *et al*. 1980; Kramer 1994; McMahon *et al*. 2003; Cutter and Ward 2005), features that might facilitate lineage-specific changes arising from directional selection on function or from diversification under either stabilizing or relaxed selection. Anecdotally, in our hands, XZ1516 was difficult to manipulate on the plate, which we hypothesize may be due to a sensitive cuticle. Moreover, another wild strain, XZ1514, was so fragile that we refrained from using it in this study, suggesting potential further genetic differentiation in collagen function across *C. elegans*.

### Comments about experimental system and design

Controlling for confounding variation poses a particular challenge in gene expression studies. For example, wild strains mature at different rates (Gems and Riddle 2000; Stastna *et al*. 2015; Zhang *et al*. 2021; Hodgkin and Doniach 1997; Poullet *et al*. 2015; Harvey and Viney 2007). We observed differences in developmental rate among our experimental strains, including that parental strain QX1211, and to a lesser extent XZ1516, its F1 with N2, and the N2 parent, developed more slowly than other strains (**Table S1**). While most F1 offspring developed at a rate similar to one parent or intermediate between the parents, the F1 offspring of QX1211 and N2 reached young adulthood over an hour faster than either parent (**Table S1**). To reduce the influence of developmental variation on gene expression differences, we harvested worms at a consistent developmental stage rather than a consistent chronological age, nevertheless all within three hours of one another (Methods). Further, we estimated the transcriptional age of each sample using an N2 gene expression time course as a ‘ruler’ (Bulteau and Francesconi 2022); all estimates fell within a five and a half hour time range (**Table S1**). These computational estimates differed across samples within strains even though such samples appeared identical and were harvested at the same time, suggesting further work is needed to understand discordance between experimental observations and computational predictions as well as inter-individual timing variation. While these measures control for developmental differences as much as is experimentally tractable, they do not guarantee the elimination of confounding developmental variation at the level of the whole organism or in tissue- or process-specific events. Thus, conclusions around specific genes should be considered carefully; for example, small differences in fertilization timing might contribute to substantial differences in the highly expressed vitellogenin genes (**Figure 6C**).

In this study, we chose the number of strains to maximize the range of genetic diversity while performing crosses simultaneously in triplicate. Maximizing the number of strains did restrict the direction of the crosses, however: we always used N2 as the maternal parent and did not include reciprocal crosses to reduce the burden of effort. The non-reciprocal cross design thus limits inferences about parent-specific transgenerational effects or sex-specific gene regulation, which have biological significance in other *Caenorhabditis* nematodes (Sanchez-Ramirez *et al*. 2021; Viswanath and Cutter 2023). For the same reason, we generated the parental strains via selfing rather than by crossing, while the F1s were obligately generated by crossing; these differences in reproductive mode may conflate with differences between F1s and parents, though we are unaware of any such broad effects on gene expression. An analytical concern with using multiple, variably diverged strains is whether ASE inferences could be standardized across strains. In theory, different strains could be differently vulnerable to reference genome bias or could exhibit increasing power to detect ASE with increasing variants within a given gene. However, the methods used here largely avoid these pitfalls (**Figure 1D, Note S1**).

Our inferences in this study, including expression classifications and trends between differently regulated genes, were robust to the inclusion or exclusion of genes in hyperdivergent haplotypes (Lee *et al*. 2021). Hyperdivergent regions differ substantially from the N2 reference sequence, making alignment and variant calling from short read data unreliable (Lee *et al*. 2021; Moya *et al*. 2025); recent RNA-seq studies in *C. elegans* sensibly and conservatively exclude genes in these regions (Lee *et al*. 2021; Zhang *et al*. 2022). However, our recent gene expression analyses showed that genome-wide trends appear robust to including or excluding genes in hyperdivergent haplotypes (Bell *et al*. 2023). Therefore, here we performed each of our genome-wide analyses both including all genes and excluding genes classified as hyperdivergent as well as genes with evidence of other possible analytical hurdles (Methods). The vast majority of trends detected when all genes were included were recapitulated when excluding hyperdivergent genes. We note, though, that results at individual genes are still likely to be influenced by genomic context, so these features should be considered when assessing small numbers of genes or conducting gene-specific queries. For example, our gene set enrichment analysis results (**Figure 2D, Figure S3, Figure S8**) were similar when including or excluding hyperdivergent genes, and whenever specific genes were used as exemplars of trends these genes were not hyperdivergent or otherwise concerning (*e.g.,* **Figure 2D**, **Figure 6C**).

In this study, we focused on global, large-scale patterns in gene level expression and did not quantify specific isoforms. However, recent evidence, and common sense, suggest that wild strains differ in expression of specific transcripts (Zhang *et al*. 2022). The extent to which non-reference strains express novel isoforms and how F1 cross progeny mediate the expression of parent-specific isoforms remain unexplored questions. A particularly intriguing possibility is that transgressive isoforms could be expressed in F1 heterozygous backgrounds but not in their native background, akin to *cis* regulatory changes that are revealed in hybrids but compensated among the parents.

### Interactive web application

Our experimental approach had many advantages, among them our study organism: the wealth of experimental data in *C. elegans* and its curation and accessibility via WormBase (Sternberg *et al*. 2024) makes this system especially amenable to analyses that add new molecular detail to existing experimental phenotypes. In turn, our characterization of gene expression variation improves our understanding of *C. elegans* and the processes that shape gene expression in this system. To aid in future genetics, trait mapping, and other *C. elegans* research, we have made the data from this study accessible via an interactive web application, where users can query their favorite gene to view its expression, regulatory pattern, inheritance mode, and other information: https://wildworm.biosci.gatech.edu/ase/.

## Data availability

Raw and processed gene expression data are available at GEO with accession number GSE272616. Per-gene per-strain data (used to perform all analyses and generate all figures), including regulatory pattern and inheritance mode classifications and underlying statistical differential expression results, are available via the Zenodo repository at https://doi.org/10.5281/zenodo.13270636. Per-gene information is interactively available via user query at web app https://wildworm.biosci.gatech.edu/ase/.

## Code availability

Code used in this study’s data processing and analysis is available at https://github.com/paabylab/wormase. Methods fully describes all existing and new software and analyses used in this study.

## Acknowledgments

We thank Samiksha Kaul and Ling Wang for help with mating plate set up and male removal during the experiment, and Han Ting Chou for undertaking a similar pilot experiment and sharing her expertise. Matt Rockman, members of the Rockman lab, Steve Burger, and three anonymous reviewers contributed helpful discussions about and suggestions for improving this work. We thank Shweta Biliya, Naima Djeddar, and Anton Bryskin at the Molecular Evolution Core Laboratory at Georgia Tech for their expert library preparation and sequencing guidance. Troy Hilley of Academic & Research Computing Services at Georgia Tech’s College of Sciences provided expert web server configuration support for the interactive web app. We gratefully acknowledge individuals in the worm community who have collected and disseminated wild *C. elegans* isolates, and the resources provided by CaeNDR (Cook *et al*. 2017; Crombie *et al*. 2024). This research was supported in part through research cyberinfrastructure resources and services provided by the Partnership for an Advanced Computing Environment (PACE) at Georgia Tech.

