## Supplemental Figures for "The regulatory architecture of gene expression variation in *C. elegans* revealed by multi-strain allele-specific analysis"

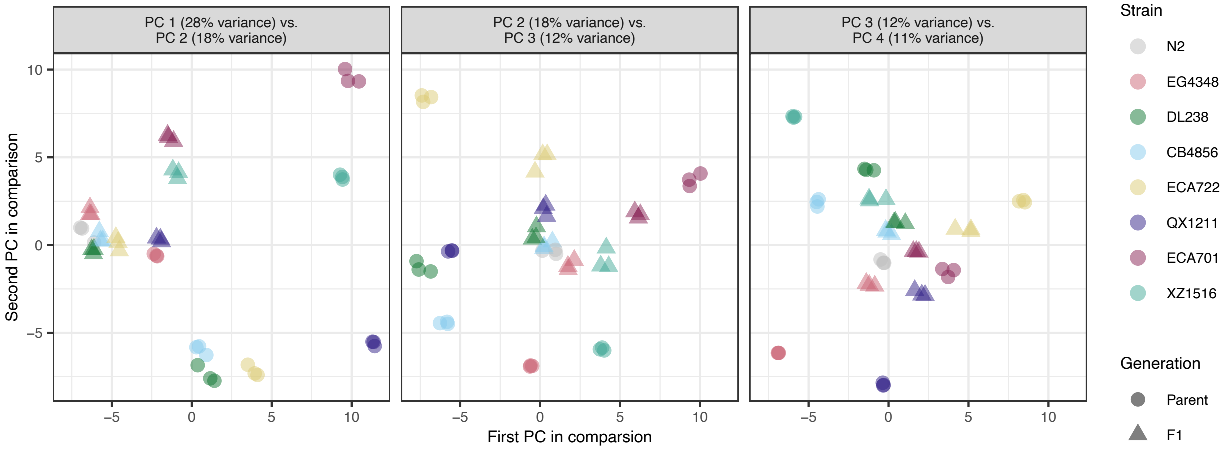


**Figure S1.** Principal components analysis (PCA) of all RNA-seq samples used in this study (*n* = 45, 3 per generation per strain). The first three pairs of principal components are plotted against one another, with percent variance explained by each component noted in the plot titles. PCA was performed using the 500 most variable genes following library size normalization, length normalization, and variance-stabilizing transformation.

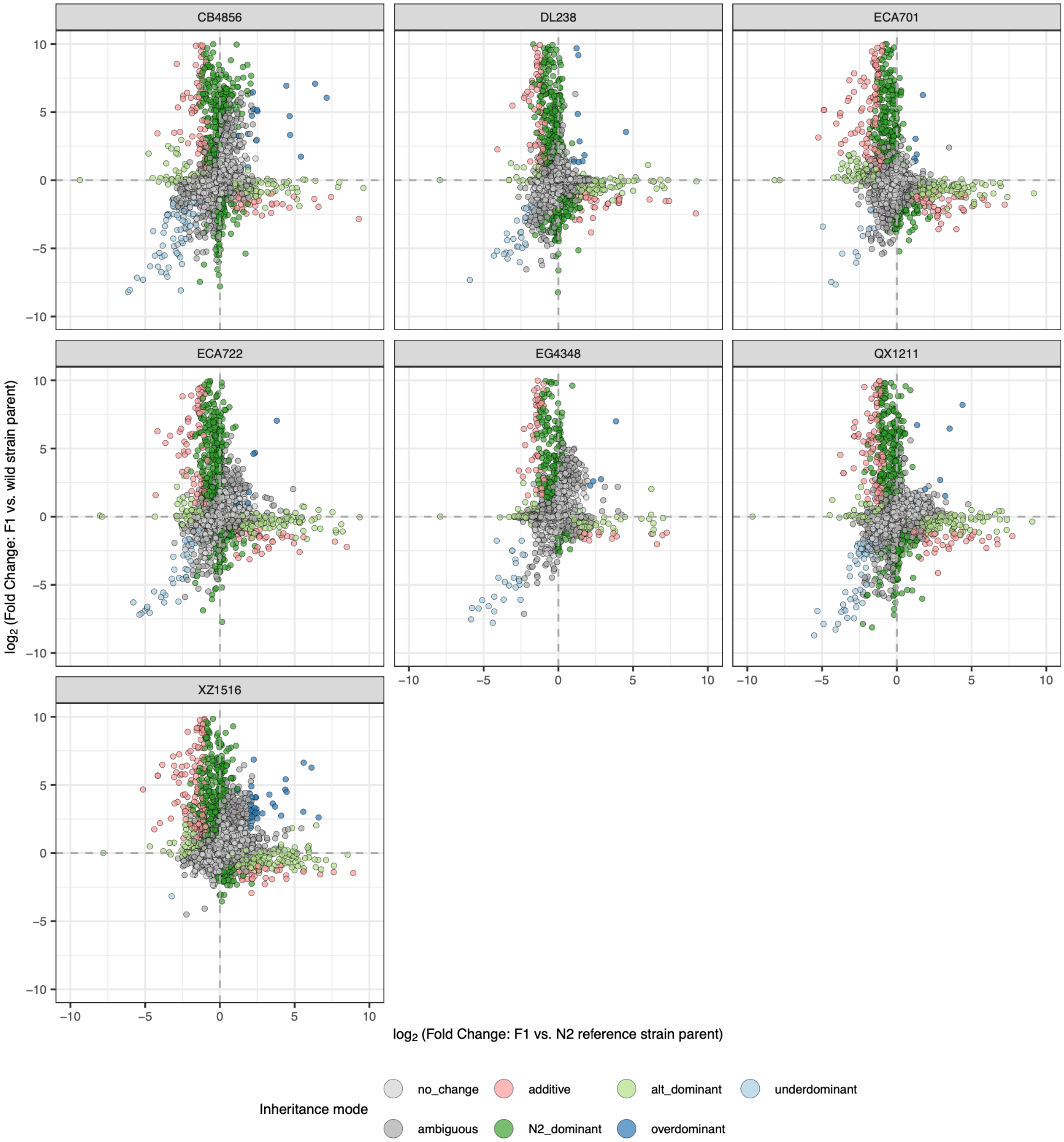


**Figure S2. Inference of inheritance mode at each gene in each strain.** As in **Figure 2A** (see Table S2 for all gene *n*s).

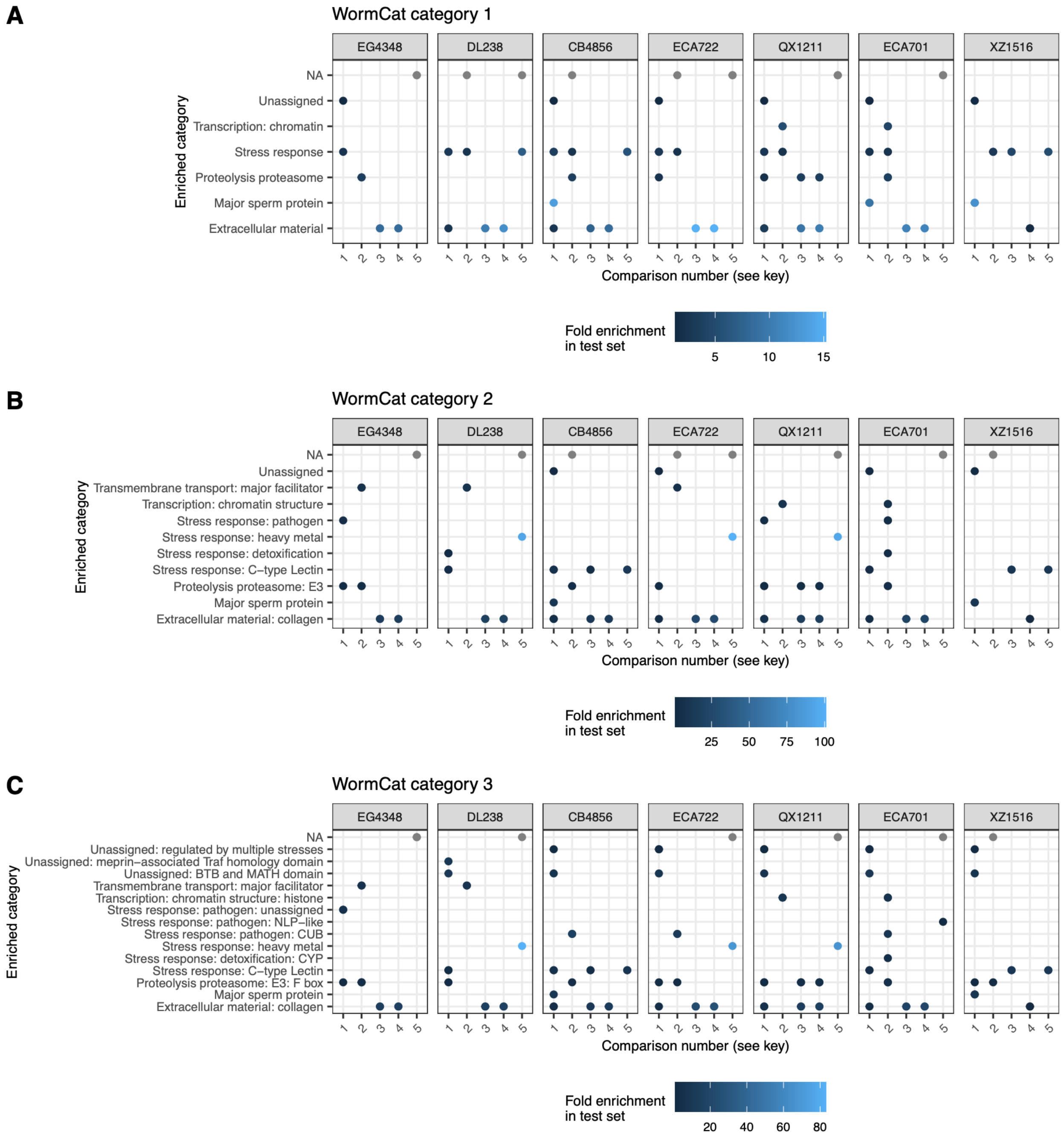


### Key

| Number | Comparison |
|---|---|
| 1 | N2_dominant vs allexpressed |
| 2 | wild_dominant vs allexpressed |
| 3 | transgressive vs allexpressed |
| 4 | transgressive_underdominant vs allexpressed |
| 5 | transgressive_overdominant vs allexpressed |

**Figure S3. All significant gene-set enrichment results for comparisons involving inheritance mode.** We performed gene set enrichment analysis for multiple comparisons (see **Key**) via a custom implementation of WormCat (Holdorf *et al.* 2020) using the appropriate test-specific background gene sets. Each category enriched in any comparison in any strain is shown: **A,** major categories (WormCat category 1); **B,** subcategories (WormCat category 2); **C,** sub-subcategories (WormCat category 3). Any blue dot indicates an enrichment at Bonferroni-corrected $p < 0.05$; the color of blue corresponds to the fold enrichment. Strains have gray dots at the dummy ‘NA’ category (top row) if that strain had no enrichments for the given comparison (to keep comparisons consistent across strains). **Figure 2C** shows differently formatted results for category 10, transgressive underdominant vs. all expressed genes.

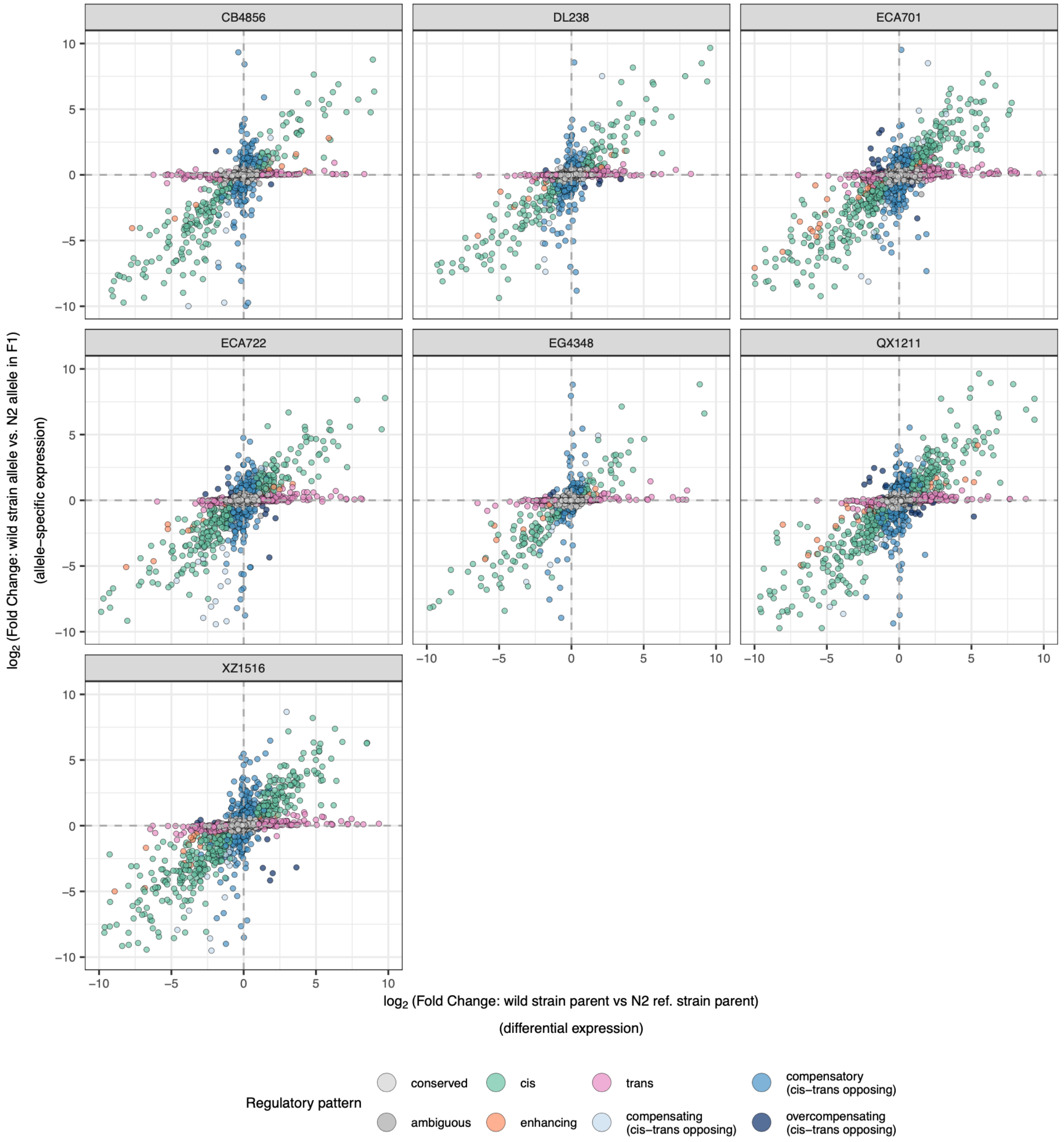


**Figure S4. Inference of regulatory pattern at each gene informative for allele-specific expression analyses.** As in **Figure 3A** (See Table S2 for all gene *n*s).

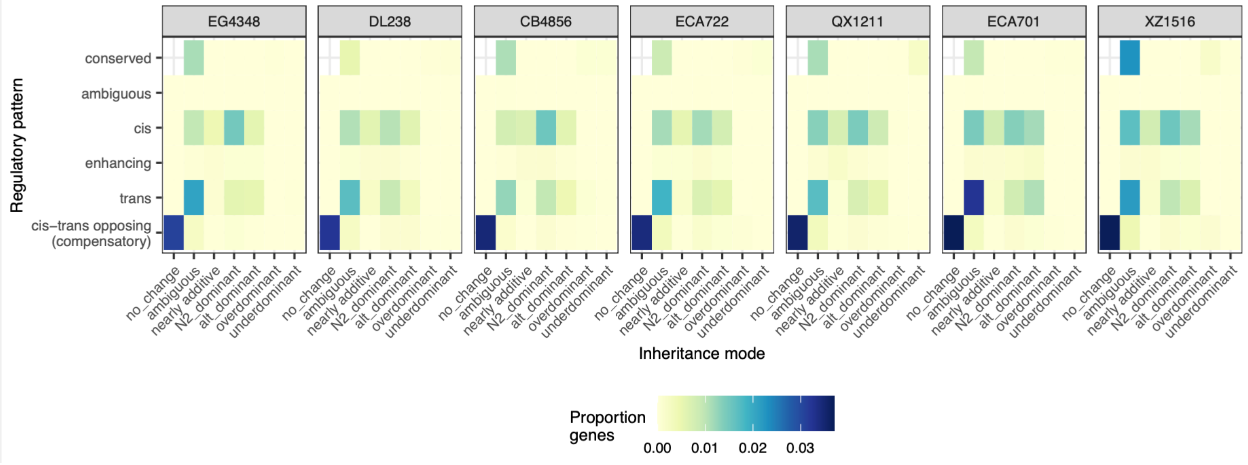


**Figure S5.** Global proportion of ASE-informative genes exhibiting each combination of inheritance mode and regulatory pattern (excluding genes without expression differences, the conserved and no change genes, for scale).

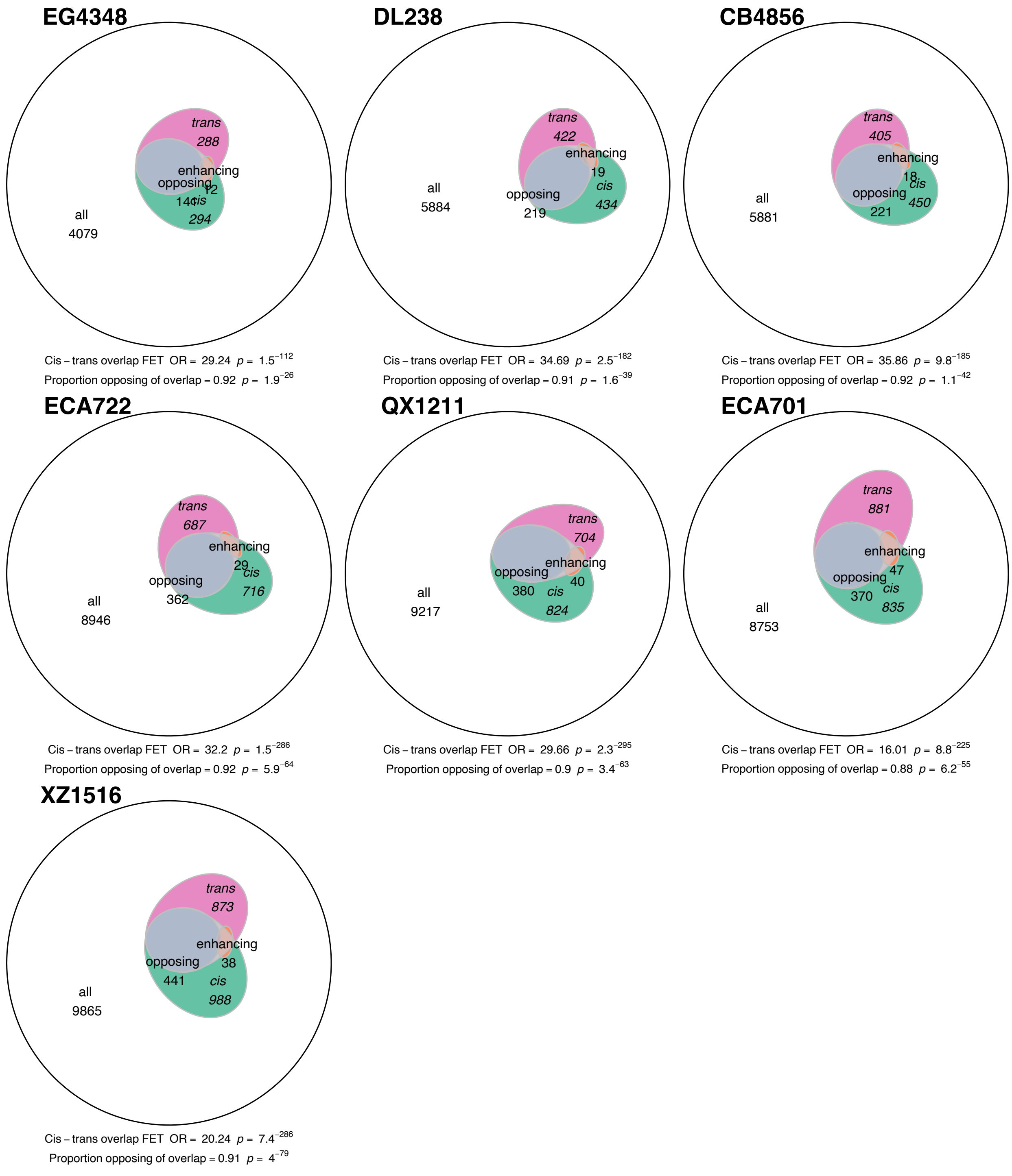


**Figure S6. Overlap of *cis* and *trans* influences on genes for all strains.** As in **Figure 3C** but for all strains. Fisher's Exact/hypergeometric test p-values for enrichment of *cis* and *trans* overlap and binomial test results for comparing the proportion of genes with overlapping *cis* and *trans* effects that oppose vs. enhance each other are shown. 'All' genes comprise all those included in this analysis, *i.e.*, allele-specific expression informative genes.

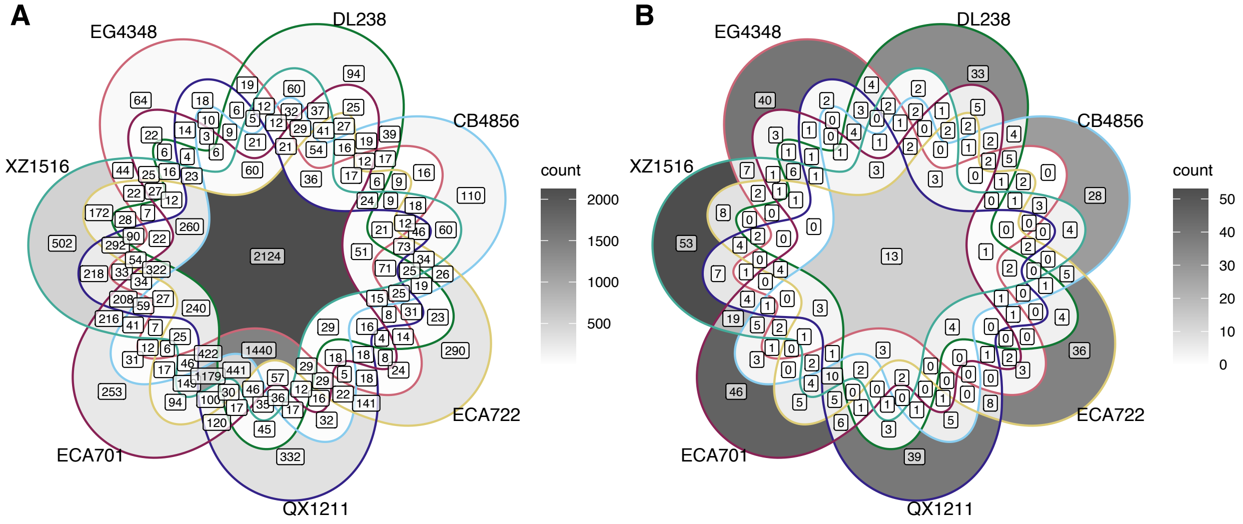


**Figure S7. Strain-wise sharing of ASE informative and allele-specifically expressed genes. A.** Genes informative for ASE in any strain are depicted in the Venn diagram based on in which strain(s) they were ASE informative. **B.** Genes that are informative across all strains (center of plot in **A**) and that have ASE in any strain are shown in the Venn diagram based on in which strain(s) they had ASE.

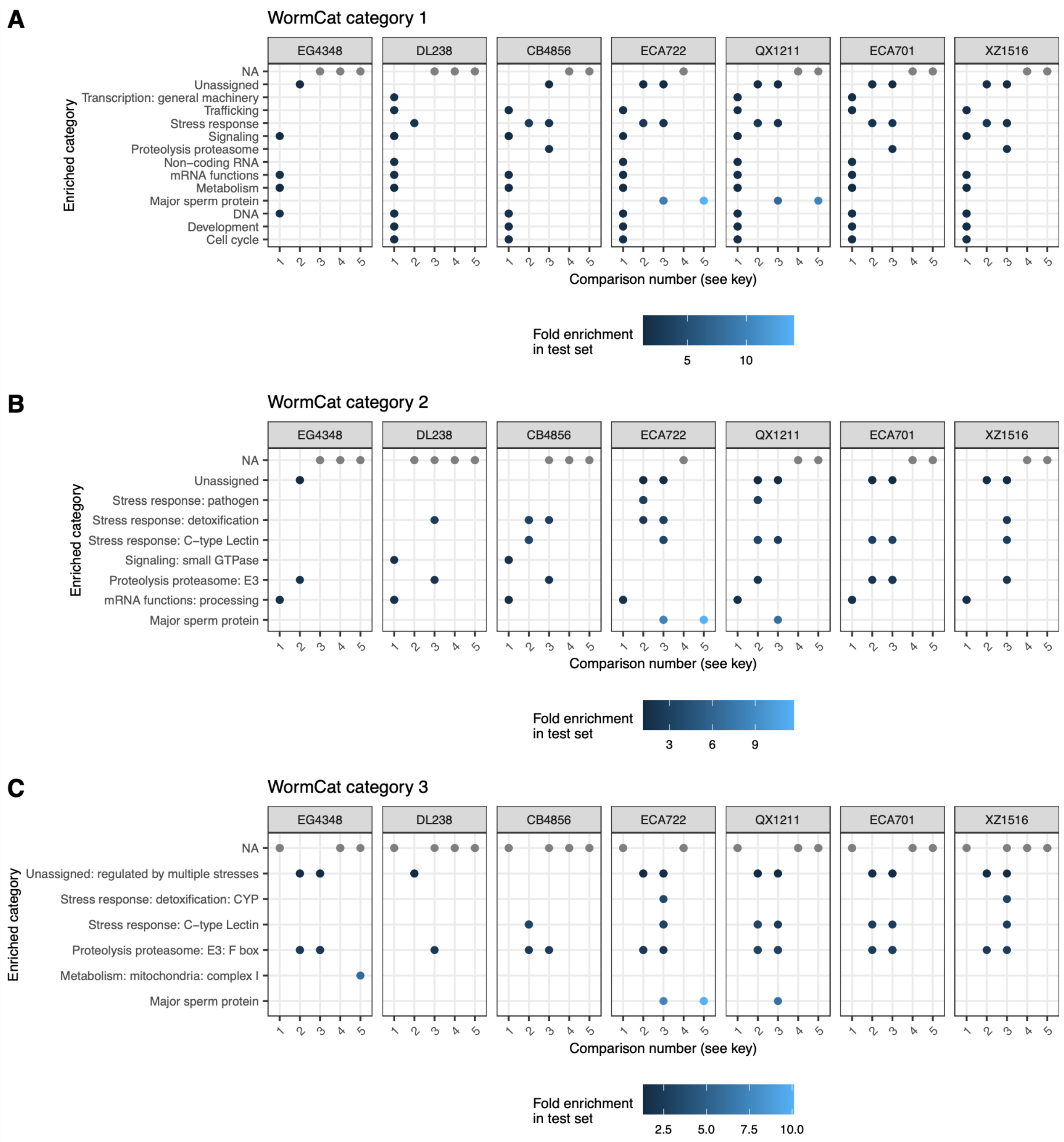


### Key

| Number | Comparison |
|---|---|
| 1 | informgenes vs allexpressed |
| 2 | degenes_inform vs informgenes |
| 3 | asegenes vs informgenes |
| 4 | compensatorygenes vs asegenes |
| 5 | compensatorygenes vs informgenes |

**Figure S8. All significant gene-set enrichment results for comparisons involving ASE and gene regulatory patterns.** We performed gene set enrichment analysis for multiple comparisons (see **Key**) via a custom implementation of WormCat (Holdorf *et al.* 2020) using the appropriate test-specific background gene sets. Each category enriched in any comparison in any strain is shown: **A,** major categories (WormCat category 1); **B,** subcategories (WormCat category 2); **C,** sub-subcategories (WormCat category 3). Any blue dot indicates an enrichment at Bonferroni-corrected $p < 0.05$; the color of blue corresponds to the fold enrichment. Strains have gray dots at the dummy 'NA' category (top row) if that strain had no enrichments for the given comparison (to keep comparisons consistent across strains).

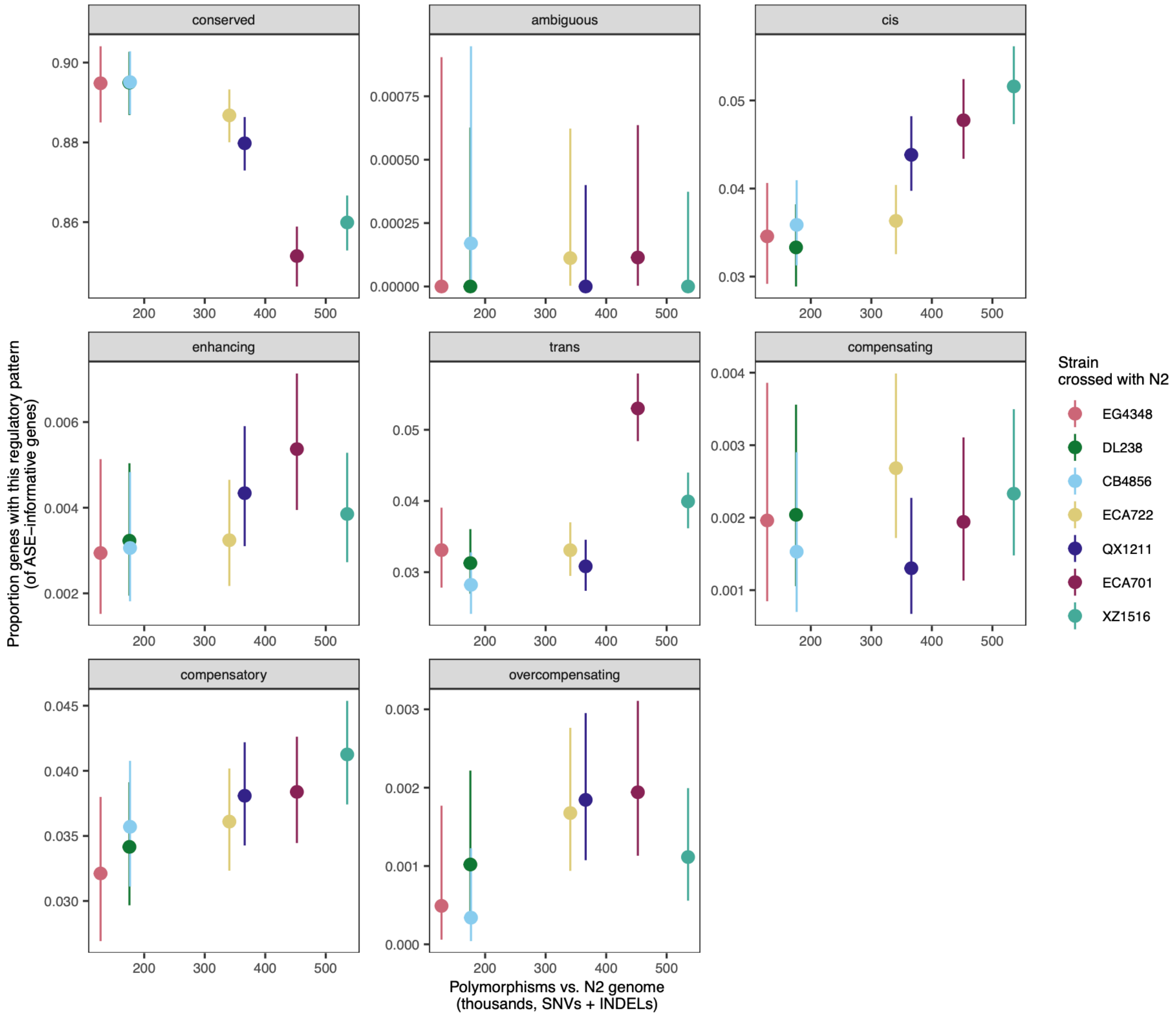


**Figure S9. Relationship between gene expression differences and genomic differentiation from N2, for each regulatory pattern category.** Genomic distance from N2 is measured as the number of polymorphisms (SNVs and INDELs) between the focal strain and the N2 reference genome. As in **Figure 4B-D** but for all regulatory pattern categories. “conserved” genes are those with conserved expression profiles across both alleles (no ASE) and strains (no DE) (all genes less the rest of the categories). The proportion of genes within each category is of all ASE informative genes in each strain (denominator differs depending on the strain) and error bars denote 95% binomial confidence intervals; all gene *n*s are provided in **Table S2**. Note that categories with relatively few genes produce less precise estimates, weakening trend inference.

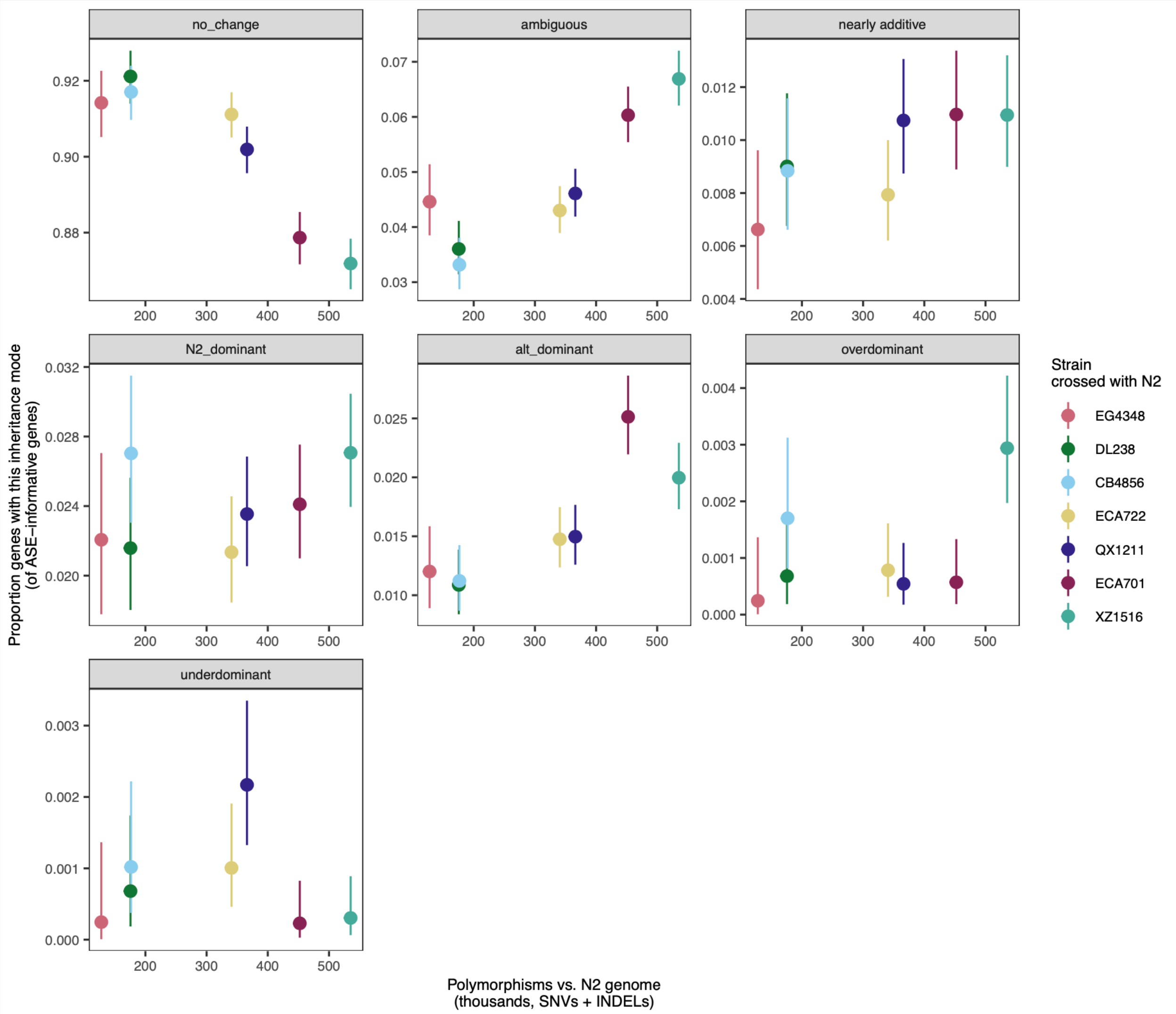


**Figure S10. Relationship between gene expression differences and genomic differentiation from N2, for each inheritance mode category.** As in **Figure 4,** but broken down into specific inheritance modes. Genomic distance from N2 is measured as the number of polymorphisms (SNVs and INDELs) between the focal strain and the N2 reference genome. "No_change" genes are those with conserved expression profiles across the parents and between the parents and F1s (all genes less the rest of the categories). Genes with wild-strain dominant expression are called "alt_dominant" here. The proportion of genes within each category is of all ASE informative genes in each strain (denominator differs depending on the strain) and error bars denote 95% binomial confidence intervals; all gene *n*s are provided in **Table S2**. Note that categories with relatively few genes produce less precise estimates, weakening trend inference.

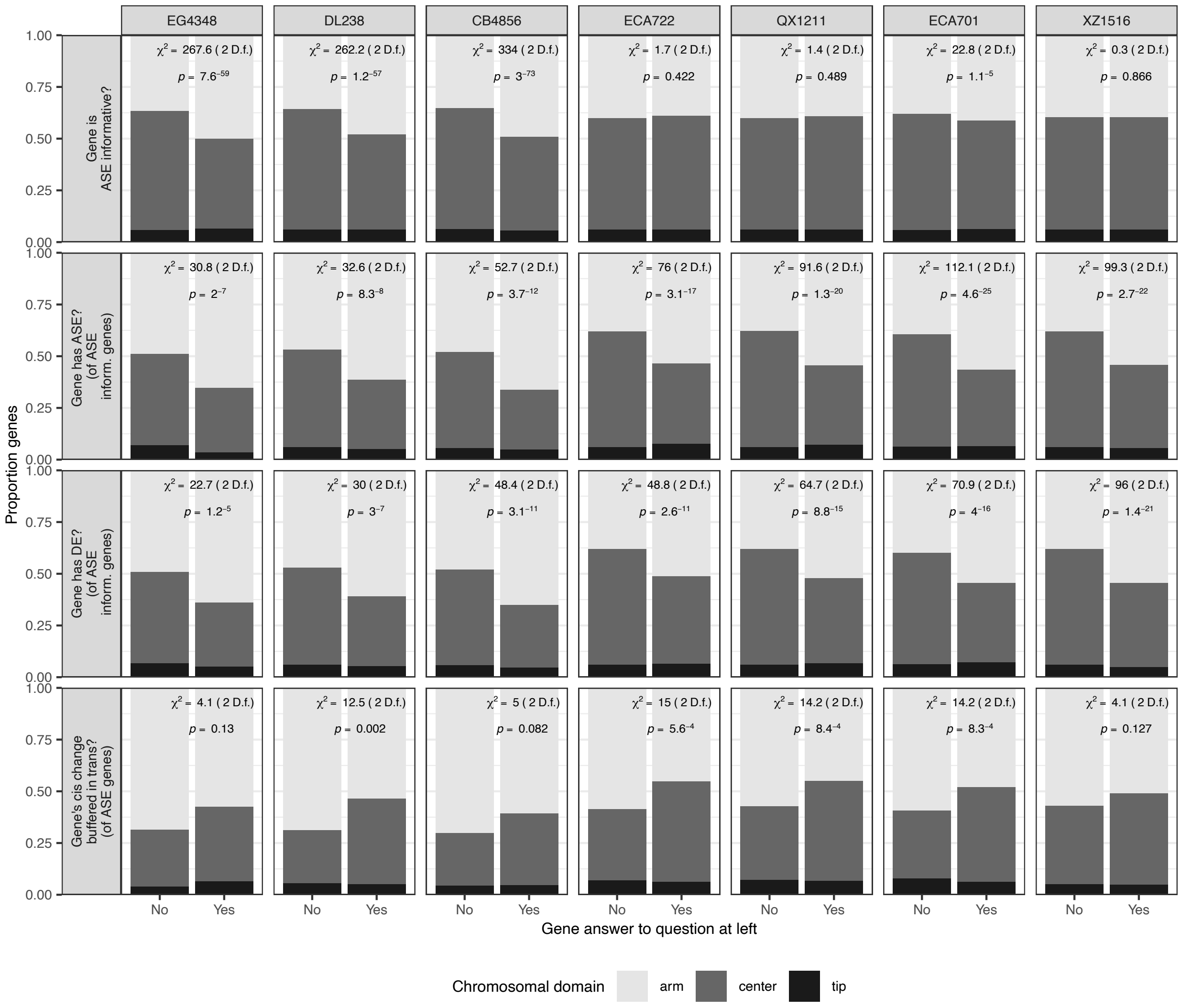


**Figure S11. Proportion of genes with expression characteristics of interest in each chromosomal domain.** Each sub-plot shows the proportion of genes that answer the question in the left plot title with the answers on the x axis broken down across chromosomal domain, separately for all strains. All underlying *n*s are in **Table S2**.
**Figure 5A** shows distribution across chromosomes for all strains combined.

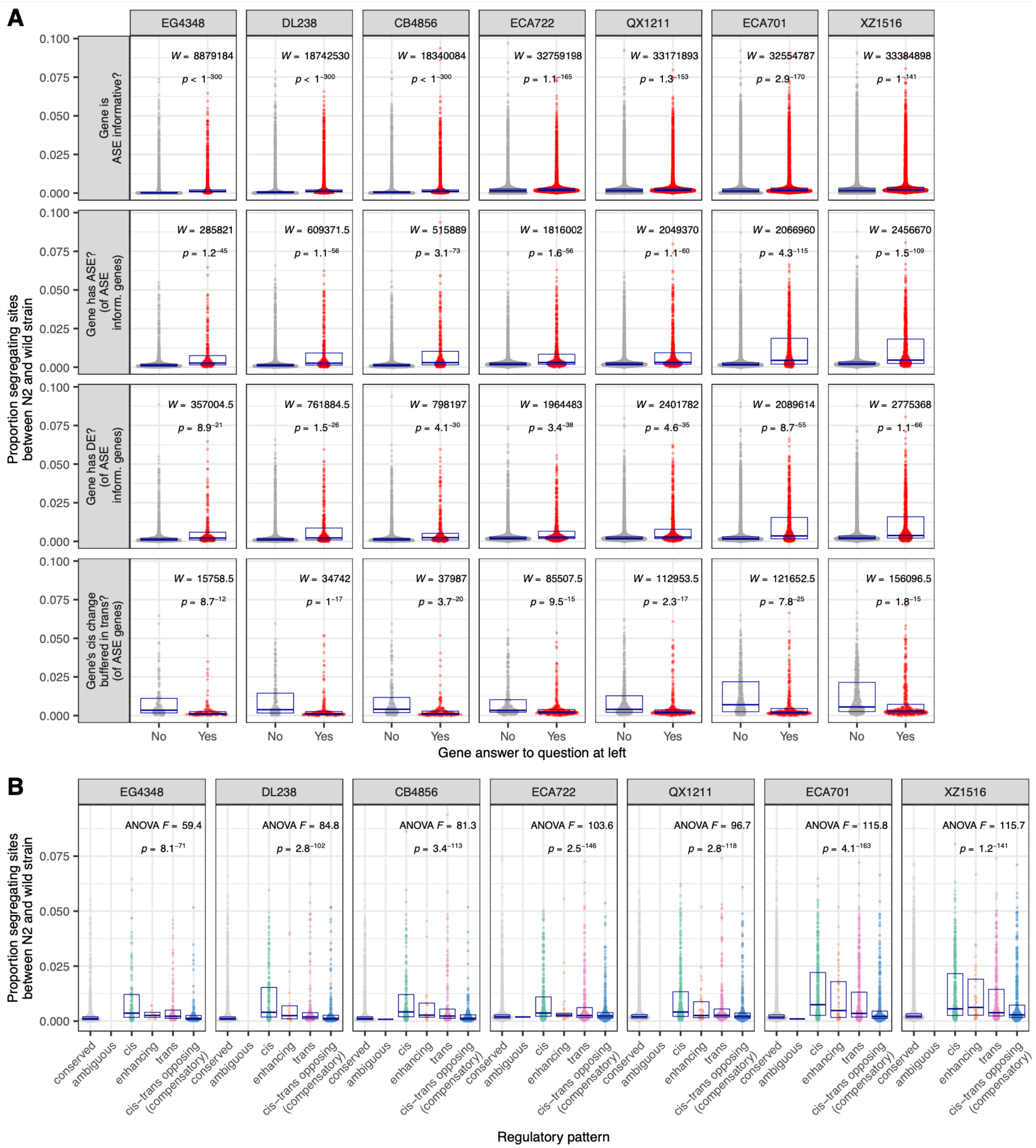


**Figure S12. Distribution of pairwise proportion segregating sites (*p*) across genes in different expression categories for each strain.** Each point represents one gene and the points fill a violin plot distribution shape; boxes denote median and interquartile range. **A.** Distribution of proportion segregating sites between the wild strain noted and N2 in genes with the x-axis label answer to the question in the left title of the plot. In bottom three rows, only ASE informative genes (middle two rows) or genes with ASE (bottom) are included, which comprise different gene sets across strains. **B.** Distribution of proportion segregating sites between the wild strain noted and N2 in each distinct regulatory pattern (called from allele-specific expression and differential expression, see text). (Of ASE informative genes; different gene sets for each strain).
All underlying *n*s are in **Table S2**.

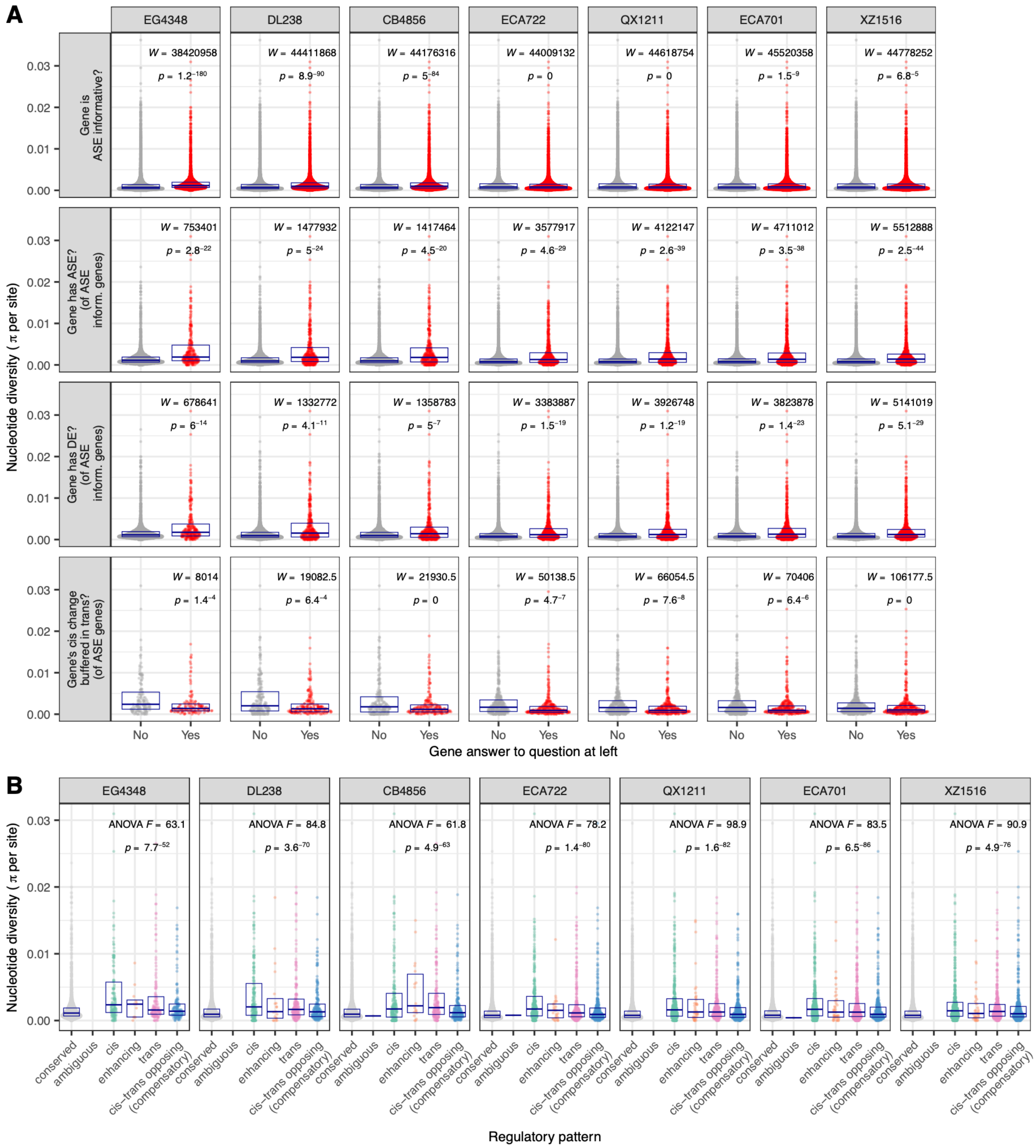


**Figure S13. Distribution of population-wide nucleotide diversity (π) across genes in different expression categories for each strain.**

**A.** As in **Figure 5B, C (left)**, but with each strain's results shown individually. In bottom three rows, only ASE informative genes (middle two rows) or genes with ASE (bottom) are included, which comprise different gene sets across strains. **B.** As in **Figure 5C (right)**. Only ASE informative genes are included, which comprise different gene sets across strains.

All underlying *n*s are in **Table S2**.

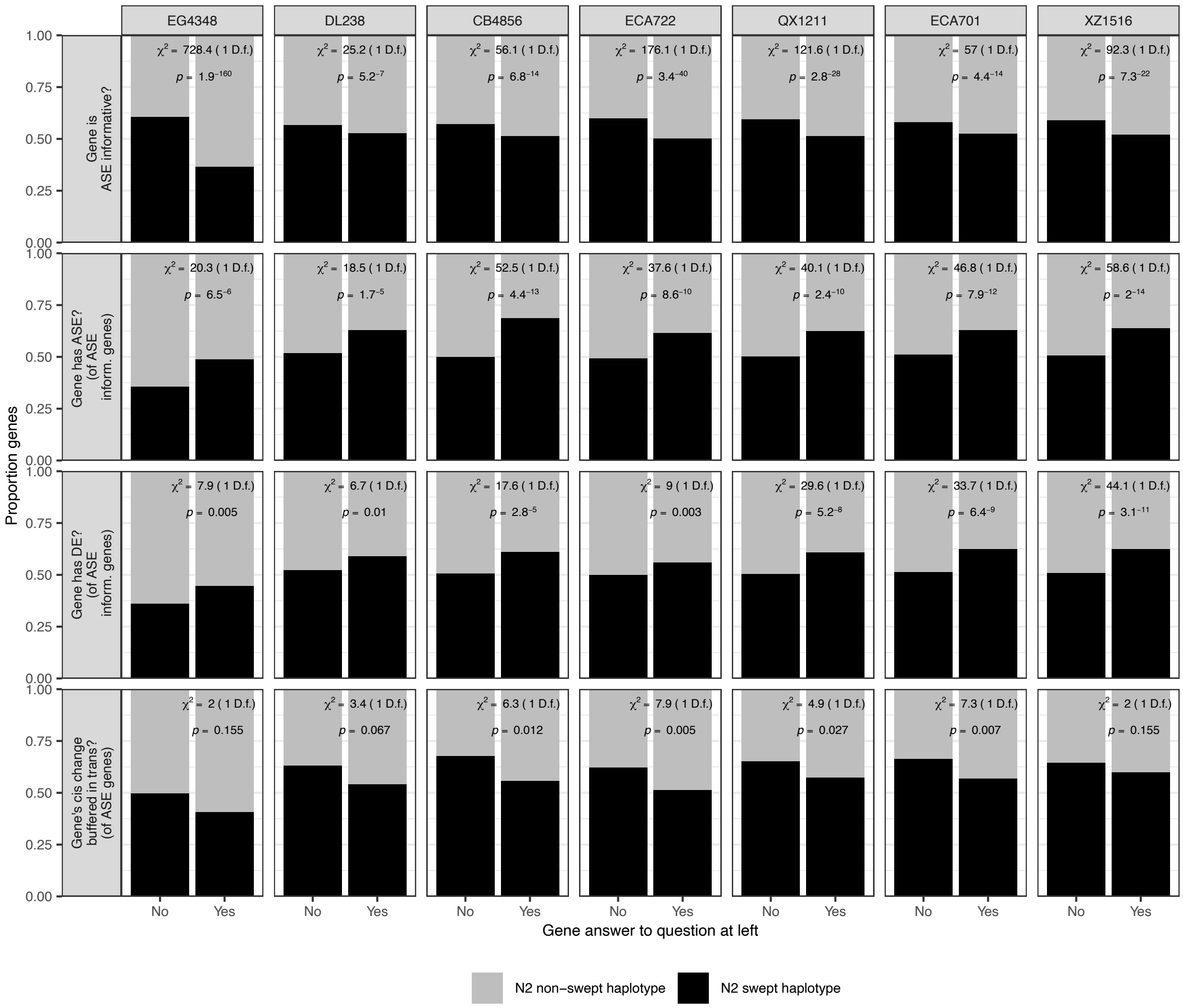


**Figure S14. Proportion of genes in each expression category of interest that are in a historically selectively swept haplotype in reference strain parent N2 for each strain.** As in **Figure 5D**, but with each strain's results individually. In bottom three rows, only ASE informative genes (middle two rows) or genes with ASE (bottom) are included, which comprise different gene sets across strains. All underlying *n*s are in **Table S2**.

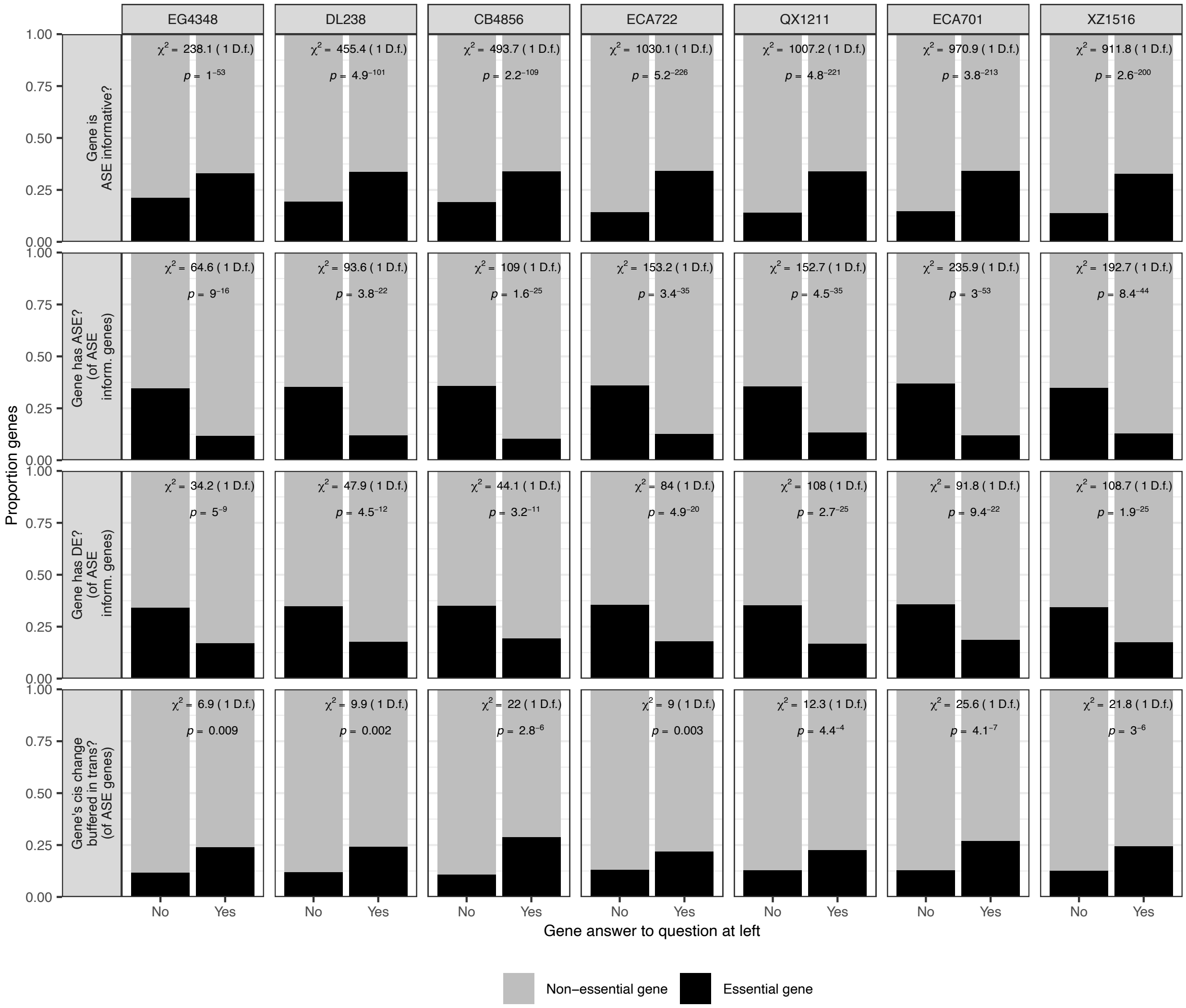


**Figure S15. Proportion of genes in each expression category of interest that are predicted essential in *C. elegans* for each strain.**

As in **Figure 5E**, but with each strain's results individually. In bottom three rows, only ASE informative genes (middle two rows) or genes with ASE (bottom) are included, which comprise different gene sets across strains. All underlying *n*s are in **Table S2**.

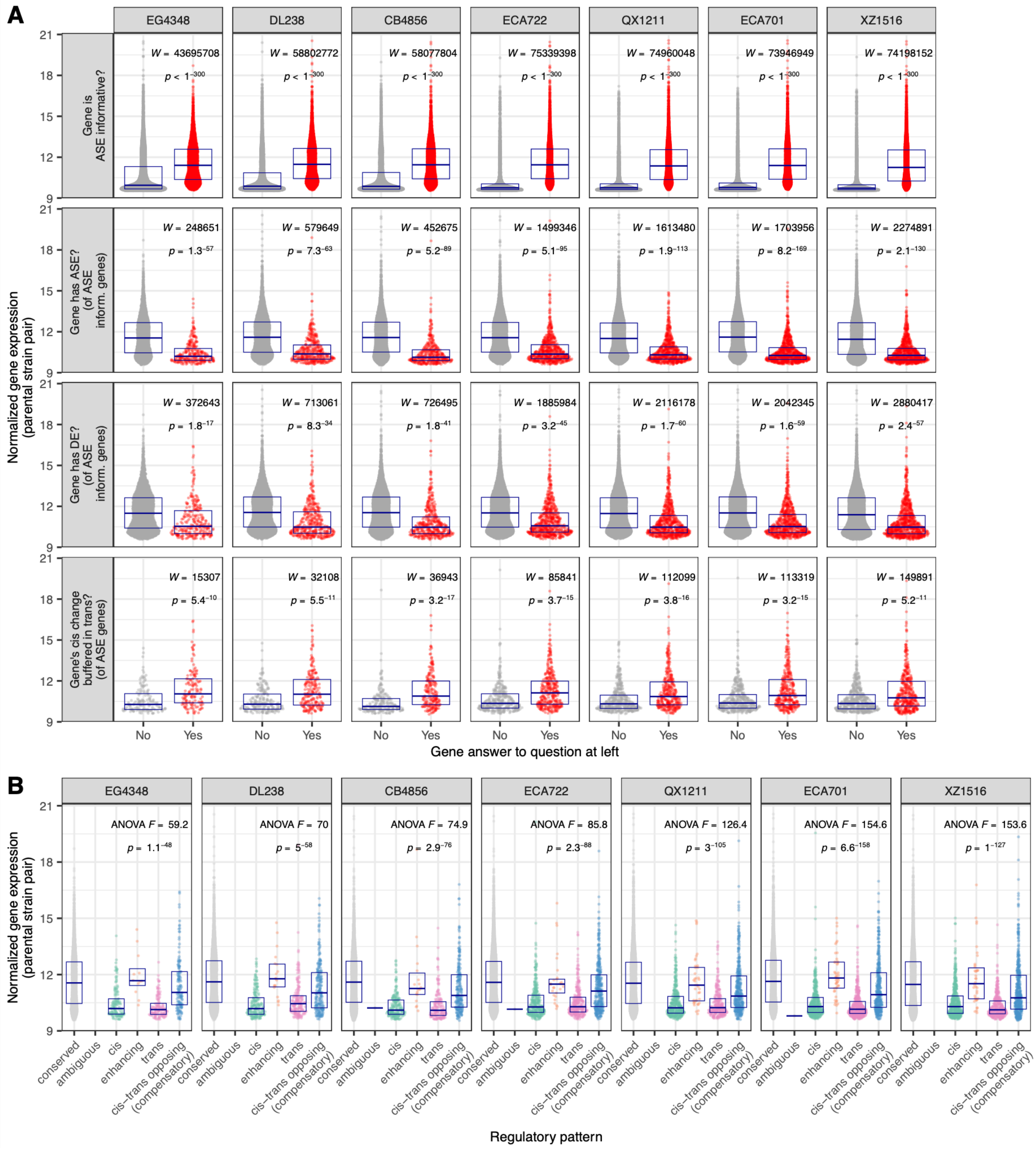


**Figure S16. Distribution of gene expression levels across genes in different expression categories for each strain.**

**A**. As in **Figure 6A, B (left)**, but with each strain's results shown individually. Each expression value is the average across samples in the parental strain for the given F1. In bottom three rows, only ASE informative genes (middle two rows) or genes with ASE (bottom) are included, which comprise different gene sets across strains. **B.** As in **Figure 6B (right)**. Each expression value is the average across samples in the parental strain for the given F1. Only ASE informative genes are included, which comprise different gene sets across strains.

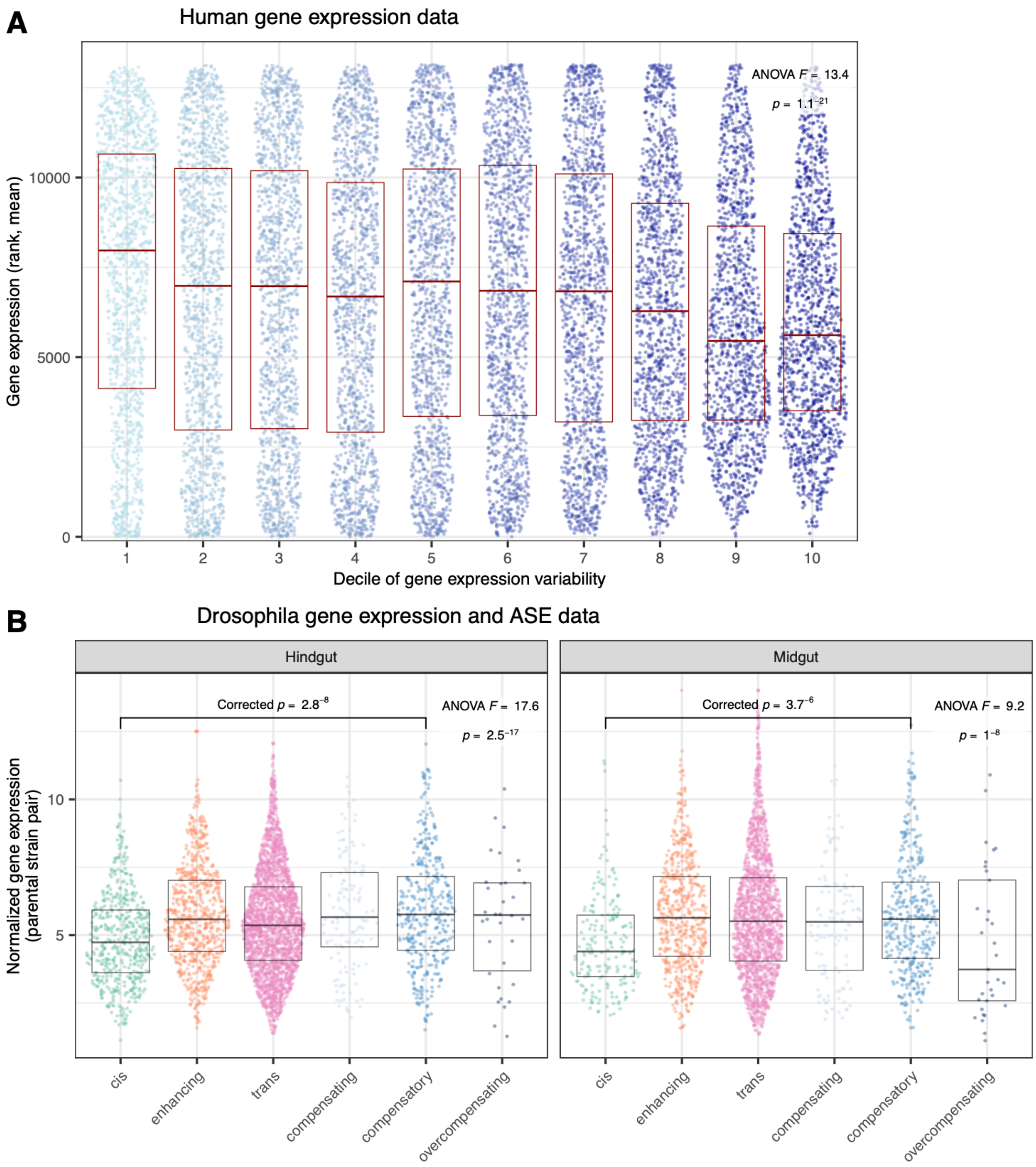


**Figure S17. Some of the observed relationships between gene expression level and gene expression variation, regulation are recapitulated in other organisms.**

**A.** Human gene expression variability *vs* gene expression level (Spearman's rho = -0.075 and $p = 8 \times 10^{-18}$). Data: rank of mean gene expression and gene's expression variance from many studies carefully combined and corrected by Wolf et al (Wolf *et al.* 2023). Each point is a gene; genes are grouped into 10 gene expression variability deciles (1: lowest 10% variability, 10: highest 10% variability) for ease of visualization; points fill a violin plot and boxes denote median +/- interquartile range. Tukey's HSD between lowest and highest variability deciles $p = 2 \times 10^{-11}$ (Bonferroni-corrected p-value; many among-decile comparisons are significant, *e.g.,* highest variability decile has significantly lower expression than 6 independent lower variability deciles) (*n* = 13,139 genes, 1313-1314 per decile). **B.** Mean normalized gene expression *vs.* regulatory pattern data from ASE experiments in four *Drosophila* crosses in two intestinal tissues (strains combined here); raw expression data and ASE and DE calls (Glaser-Schmitt *et al.* 2024). Each point represents one gene in one strain; Y axis denotes gene expression amount (length, library size, and variance-log2 normalized, Methods). Here, only genes with changed regulation are shown to minimize differences among studies in including informative genes in analyses. Tukey's HSD comparisons (Bonferroni multiple testing adjusted) for compensatory *vs* cis-dominant regulated genes are shown on the plot; other comparisons were also significant (notably, trans and enhancing > cis). *N* genes = 4857 hindgut, 4282 midgut; each is plotted four times, once for each strain comparison. Box plots show median and 1.5x IQR.
