## Supplementary material for "The regulatory architecture of gene expression variation in *C. elegans* revealed by multi-strain allele-specific analysis": Note S1

### Supplementary Note 1 (Note S1)
### Investigating if increased genetic variation within informative genes introduces differences in power for calling ASE

Performing an allele-specific expression (ASE) study across multiple strain pairs with varying levels of divergence from one another enables important analyses about the relationship between divergence and expression variation and evolution (see main text). However, one concern is that the degree of divergence might affect the power to detect ASE. For example, more diverged strains might suffer from greater reference bias (wherein their more-diverged sequences fail to map to the reference genome), resulting in artifactual calls of genes with reference-skewed ASE. However, our method of making strain-specific genomes followed by careful quantitation, allelic estimation, and significance testing minimizes concerns about reference bias (**Figure 1D**).

Alternatively, it is possible that the increased variation among more diverged strains could make ASE detection easier: the increase in variants could make alignment to the correct haplotype easier, vs. aligning non-specifically to both haplotypes, resulting in more power to detect ASE and thus the detection of more ASE in F1 offspring from more-diverged strain pairs. Requiring each gene examined for ASE to have at least 5 haplotype-and-gene specific alignments—creating the ASE-informative gene set—somewhat allays this concern, but doesn't account for potential differences in strains with, e.g., 5 specific alignments at a gene vs. 10 specific alignments at a gene due to more variants within the coding region. Comfortingly, the among-strain patterns we observed (main text) were robust to different thresholds for ASE-informative genes (at least 2, 5, and 10 haplotype-specific alignments). However, we still sought to further understand if ASE detection might remain more powered for more diverged strain pairs.

First, we determined that, as expected, higher-diverged strain pairs did have, on average, lower proportions of non-haplotype-specific alignments (**Figure SN1.1**). In our workflow, the number of haplotype-specific alignments was determined from the output of Salmon (Patro *et al.* 2017) pseudomapping RNA quantitation: Salmon constructs "equivalence classes" for each set of similar reads; these equivalence classes can be shared across isoforms, genes, and alleles, or unique to those categories. In our workflow, these equivalence classes are used both to define which genes are informative (those with 5+ haplotype-specific alignments in each sample of a strain) and as input to EMASE (Raghupathy *et al.* 2018), which uses them and a maximum-likelihood approach to estimate final per-allele and overall counts. These EMASE estimated counts are what is used for ASE significance testing (by DESeq2 (Love *et al.* 2014)). Therefore, the only information we have on the uniqueness of specific alignments comes from data that resides multiple downstream steps away from actual ASE testing, so it is unclear if the relationship between divergence and unique alignments (**Figure SN1.1B**) has any downstream impact on ASE testing and power to call ASE.

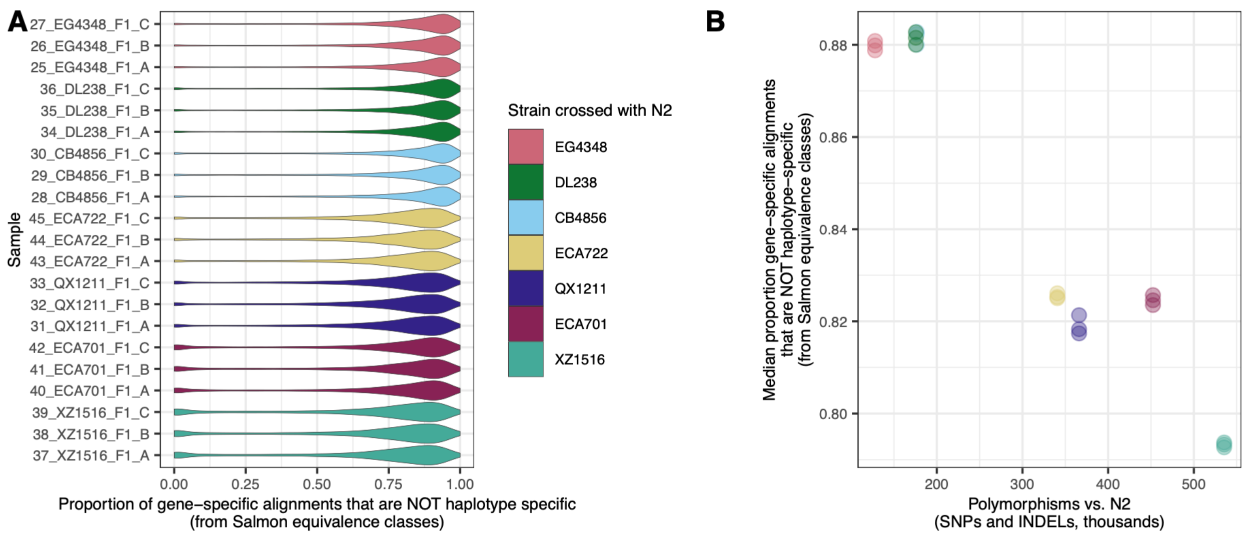


**Figure SN1.1.** Per-gene distribution of alignments that are not haplotype specific in each sample (**A**) and median gene's proportion non-specific alignments plotted against divergence from the reference strain (**B**); colors are consistent across panels. All informative genes were included in the analysis (ns in Table S2). The underlying numbers come from analyzing Salmon (Patro *et al.* 2017) equivalence classes for which equivalence classes are specific to a given haplotype and gene, and then computing the proportion of the total gene's alignments that are assigned to these equivalence classes. Note that these data are multiple steps upstream from ASE statistical testing. In (**B**), each sample is one point and the CB4856 samples (light blue) are obscured by the DL238 samples (dark green).

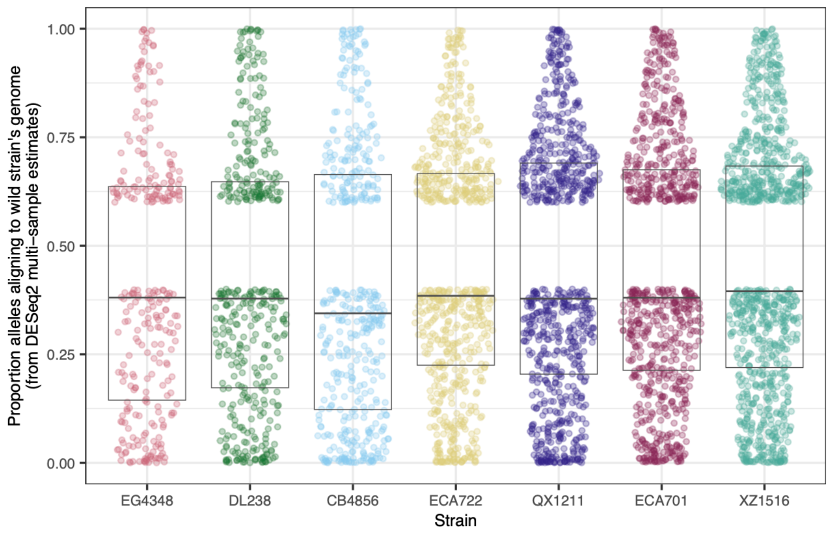


**Figure SN1.2.** Proportion of alleles assigned to the wild strain after final ASE processing at each gene with ASE called (by DESeq2 (Love *et al.* 2014); after EMASE (Raghupathy *et al.* 2018) processing of the Salmon allele counts in **Figure SN1.1**). Data is the same as shown in **Figure 1D** (main text), but showing only each gene with ASE called (ns in Table S2). Strains are ordered according to divergence from N2; box plots show median and interquartile range.

So, we sought to examine whether the estimates used for ASE detection itself show any bias based on the divergence of parental genomes from one another; **Figure 1D** (main text) shows this for all genes. Here, we narrow specifically to genes with ASE called: is there any detectable bias in the estimated allelic proportion based on divergence (**Figure SN1.2**)? There does not seem to be. While these data are much more relevant to actual ASE calls (they are generated by combining EMASE counts across samples and modeling by DESeq2), they are not integrated with the specific unique alignments data, so, while comforting, this analysis can't directly test the overall concern.

Another way to try to understand whether this more-uniqueness-with-divergence is a big problem in our framework is to look at the range of haplotype-unique mapping across strains (**Figure SN1.3**). We imagine that if more haplotype-uniqueness (with divergence) led to higher power to detect ASE, genes would be more likely to be called as having ASE in all the strains if they were more similar in the haplotype-unique proportion across strains. Put another way, we'd expect a large range of haplotype-uniqueness across strains to result in ASE being called in some strains (presumably those with more haplotype-unique alignments) than in others. We do not see this trend: of genes that are informative for ASE analyses in all strains, genes called ASE in only one strain had on average a lower range of haplotype-unique alignment proportion than genes called ASE in multiple strains (**Figure SN1.3**). Genes in which six strains (all but one) had ASE called tended to have a wider range in unique alignments, but this range did not preclude them from being detected as ASE in these six strains (**Figure SN1.3**). We think this result confers reasonable confidence that overall, our ASE detection power is not strongly biased toward more diverged strains.

Still, this is a broad-strokes approach and a tricky issue. To try to develop a better understanding, we looked at the proportion of haplotype-unique alignments, divergence, and ASE calls at the same time for some example genes (**Figure SN1.4**). First, we examined *fog-2* (**Figure SN1.4A**), the gene that was knocked out in the N2 parent to require obligate outcrossing. Sensibly, all strains had significant ASE skewed toward the wild strain parent at this gene (90-93% of alleles called as wild strain by DESeq2): the wild strain copy of *fog-2* was theoretically the only one being expressed. While more diverged strains did have lower proportions of non-unique alignments here, all strains had ASE called and at similar magnitude (**Figure SN1.4A**). To see if this trend extended to genes with lower allelic skews, closer to our calling thresholds, we looked for genes with close to 60% of one allele or the other and called ASE in one or more strains. One relatively highly expressed gene was *clp-7* (**Figure SN1.4B**); it had ASE called in two strains but not the remaining three – but strains with both high and low haplotype-mapping proportions were called both as ASE and as not ASE, suggesting that haplotype-specific mapping was not a major factor in ASE determination in this case. A lower expressed gene with some ASE estimates close to the 60% threshold was *fkb-8* (**Figure SN1.4C**); again here, some strains were called ASE with both high and low proportions of haplotype-specific alignments. While obviously not exhaustive, analysis of these example genes suggests that the proportion of Salmon equivalence class alignments that map only to one haplotype is not a definitive factor in calling ASE.

While we cannot entirely rule out higher power to detect ASE in more diverged strain pairs, taken together, the lines of evidence presented here suggest that power to call ASE is not strongly biased towards strains with more divergence within genes already thresholded as informative for ASE analyses.

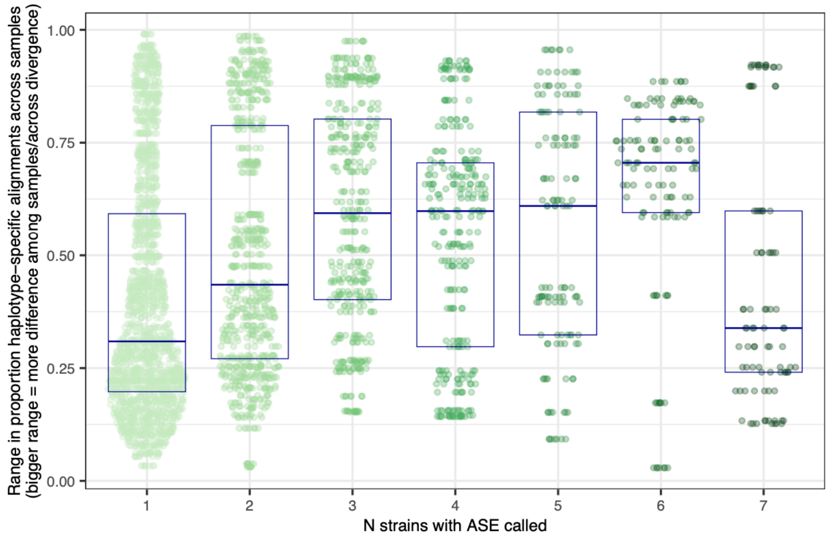


**Figure SN1.3.** The range in samples' proportion of haplotype-specific alignments (y axis) for each gene called ASE in one or more strains (x axis), of genes ASE informative in all strains. A range of 0 means that all samples have the same proportion of alignments mapping haplotype-uniquely at that gene, while a range of 1 means that one sample has no uniquely mapping alignments at that gene while another sample has all alignments at that gene uniquely mapping.

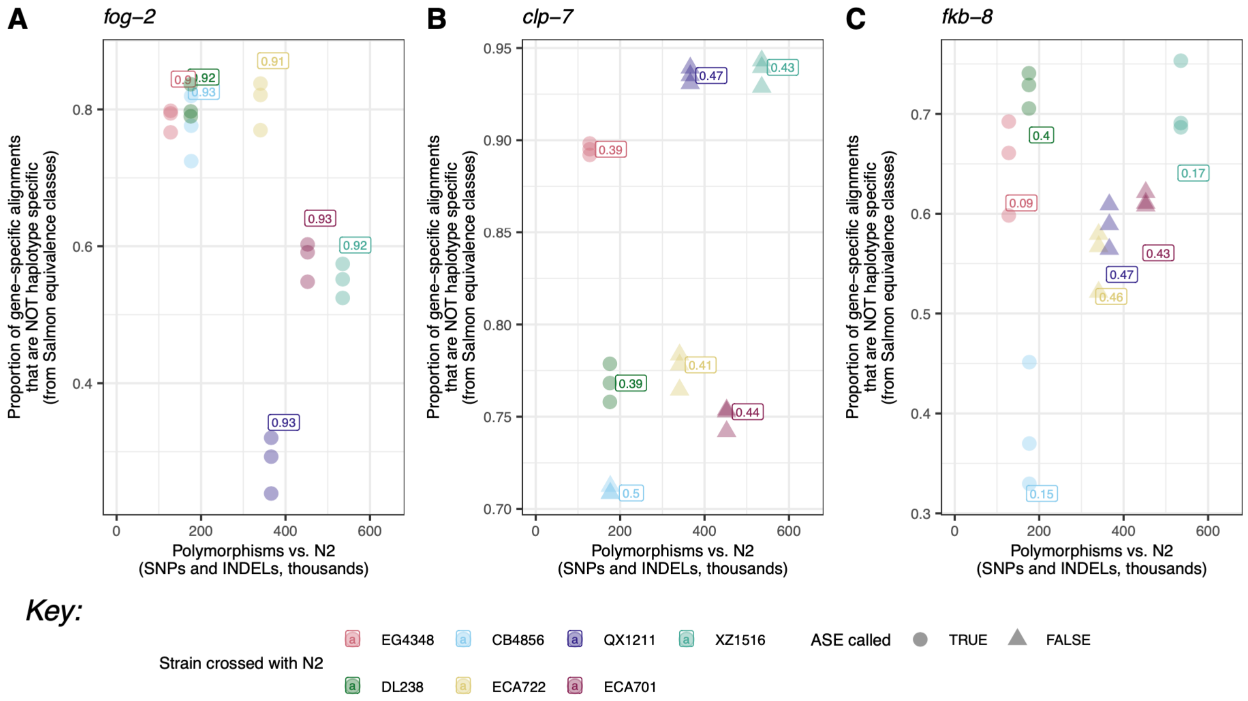


**Figure SN1.4.** Examples of genes called ASE in at least some strains (point shape denotes if ASE was called in that strain) with the proportion of alignments at that gene that do not uniquely map in each sample (y axis). Each point is one sample ($n$ = 21 F1 samples, 3 per strain) and each group of points obligately has the same point shape, as ASE is called on a per-strain rather than per-sample basis. The numbers annotating each group of points are the proportion of wild-strain alleles called by DESeq2 for ASE testing in that strain (same data as y axis **Figure SN1.2**). Note that higher non-uniquely mapping proportions (higher y-axis) do not necessarily lead to failure to call ASE, while higher non-uniquely mapping proportions (lower y-axis) do not preclude ASE calling. **A** shows *fog-2*, the gene knocked out in the N2 parent to force N2 to outcross rather than self-mate. **B** shows *clp-7*, a gene with ASE called but allelic proportion close to the threshold 60% of one allele in at least one strain and relatively high expression level. **C** shows *fkb-8*, a gene with ASE called but allelic proportion close to the threshold 60% of one allele in at least one strain and relatively low expression level.

**Note S1 References**

Love MI, Huber W, Anders S. Moderated estimation of fold change and dispersion for RNA-seq data with DESeq2. *Genome Biol* 2014;15(12):550. 10.1186/s13059-014-0550-8

Patro R, Duggal G, Love MI, Irizarry RA, Kingsford C. Salmon provides fast and bias-aware quantification of transcript expression. *Nat Methods* 2017;14(4):417-419. 10.1038/nmeth.4197

Raghupathy N, Choi K, Vincent MJ, Beane GL, Sheppard KS, Munger SC, Korstanje R, Pardo-Manual de Villena F, Churchill GA. Hierarchical analysis of RNA-seq reads improves the accuracy of allele-specific expression. *Bioinformatics* 2018;34(13):2177-2184. 10.1093/bioinformatics/bty078
