## Supplementary material for "The regulatory architecture of gene expression variation in *C. elegans* revealed by multi-strain allele-specific analysis": Note S2

### Supplementary Note 2 (Note S2)

### Avoiding artifactual *cis-trans* estimate negative correlation and commensurate inflation of inferences of cis-trans compensation

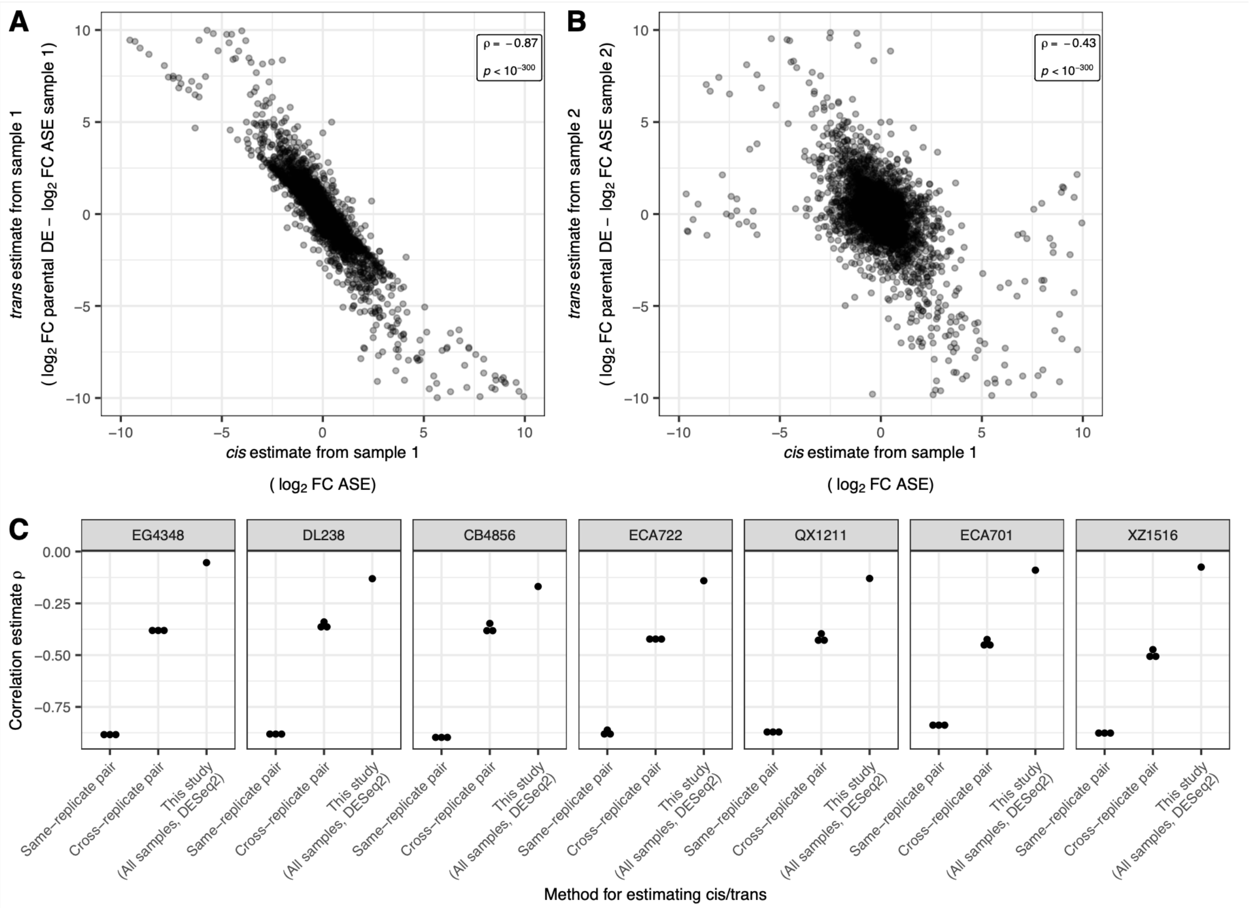


**Figure Supplementary Note 2.1 (SN2.1). An explanation of artifactual *cis-trans* estimates' autocorrelation and how our method nullifies this artifact.**
**A**. Negatively correlated *cis*-*trans* estimates from using one ECA722 strain sample to estimate the ASE used to determine both *cis* and *trans* estimates. *n* genes = 8947 (ASE informative in this strain; 89 ommitted from plot due to exceeding axis limits) **B.** Decreased negative correlation of *cis*-trans estimates using cross-replicate correction (Fraser 2019), where one sample is used to estimate the extent of *cis* (x axis) and a different sample was used to generate the ASE estimate used to estimate the extent of *trans* (y axis). *n* genes = 8947 (ASE informative in this strain; 91 omitted from plot due to exceeding axis limits). **C.** Correlation coefficients (Spearman's $\rho$) between *cis* and *trans* estimates for multiple ways of estimating *cis* and *trans* for each studied strain. When using pairs of replicates, there are three possible pairs because each strain was sequenced in triplicate, and therefore three correlation coefficients are shown. When using this study's method, all samples are used, so the single correlation coefficient estimate is shown. Less negative correlations suggest less artifactually-introduced correlation.

The extent to which a gene is regulated in *cis* is estimated as the extent of allele-specific expression at that gene because the *trans* environment is shared across the alleles in a hybrid individual (Cowles *et al.* 2002; Yan *et al.* 2002). On the other hand, differential expression between strains arises from the combination of of *cis*- and *trans*-acting forces (Wittkopp *et al.* 2004). Therefore, the amount of *trans* regulation is estimated as the total between-strain differential expression minus the extent of *cis* difference (ASE). When ASE is estimated once, for example in one sample, this same estimate is used to derive both total *cis* estimate and the total *trans* estimate, so these *cis* and *trans* estimates being automatically negatively correlated (Fraser 2019; Zhang and Emerson 2019) (**Figure SN2.1A**). This relationship means that if an in the case of a spurious inference of ASE, the gene will be called compensatory due to lack of differential expression; this effect over-inflates estimates of *trans* compensation of *cis* changes (Fraser 2019; Zhang and Emerson 2019).

Fraser (2019) recommends avoiding this issue via "cross-replicate correction:" by using one sample to estimate the ASE used to define *cis* and a different sample to estimate the ASE used to define the extent of *trans* regulation. Similarly, Emerson and colleagues (Emerson *et al.* 2010; Zhang and Emerson 2019) avoid bias by using different technical replicates to estimate the ASE used for *cis* and *trans* estimates. We found that cross-replicate correction indeed reduced the negative correlation of *cis* and *trans* when compared with same-replicate comparisons in our data (**Figure SN2.1B,C**).

We reasoned that our analytical approach of modeling ASE using multiple samples at once might also negate the artifactual *cis-trans* autocorrelation; indeed, our multi-sample derived estimates of ASE result in a markedly less negative correlation between estimates of *cis* and *trans* than even the cross-replicate correction (**Figure SN2.1C**). Furthermore, instead of taking our numerical estimates of ASE and DE as true point estimates of the extent of *cis* and *trans*, we instead use them to categorize genes into regulatory pattern categories, which further insulates our inferences from biases introduced by incorrect estimations and their autocorrelation.

*Specific definitions/methods*
ASE: log2(fold change wild allele / N2 allele)
DE: log2(fold change

Within sample (for purposes of this investigation only): EMASE (Raghupathy *et al.* 2018) counts used directly to estimate ASE within each sample; DE was still estimated using multiple samples and DESeq2 (Love *et al.* 2014) modeling. See Methods for modeling used for our full analyses.

**Note S2 References**

Cowles CR, Hirschhorn JN, Altshuler D, Lander ES. Detection of regulatory variation in mouse genes. *Nat Genet* 2002;32(3):432-437. 10.1038/ng992

Emerson JJ, Hsieh LC, Sung HM, Wang TY, Huang CJ, Lu HH, Lu MY, Wu SH, Li WH. Natural selection on cis and trans regulation in yeasts. *Genome Res* 2010;20(6):826-836. 10.1101/gr.101576.109

Fraser HB. Improving Estimates of Compensatory cis-trans Regulatory Divergence. *Trends Genet* 2019;35(1):88. 10.1016/j.tig.2018.10.003

Love MI, Huber W, Anders S. Moderated estimation of fold change and dispersion for RNA-seq data with DESeq2. *Genome Biol* 2014;15(12):550. 10.1186/s13059-014-0550-8

Raghupathy N, Choi K, Vincent MJ, Beane GL, Sheppard KS, Munger SC, Korstanje R, Pardo-Manual de Villena F, Churchill GA. Hierarchical analysis of RNA-seq reads improves the accuracy of allele-specific expression. *Bioinformatics* 2018;34(13):2177-2184. 10.1093/bioinformatics/bty078

Wittkopp PJ, Haerum BK, Clark AG. Evolutionary changes in cis and trans gene regulation. *Nature* 2004;430(6995):85-88. 10.1038/nature02698

Yan H, Yuan W, Velculescu VE, Vogelstein B, Kinzler KW. Allelic variation in human gene expression. *Science* 2002;297(5584):1143. 10.1126/science.1072545

Zhang X, Emerson JJ. Inferring Compensatory Evolution of cis- and trans-Regulatory Variation. *Trends Genet* 2019;35(1):1-3. 10.1016/j.tig.2018.11.003
